# AART enables fast and accurate cross-platform proteomic translation

**DOI:** 10.64898/2026.06.29.735313

**Authors:** Yurui Chen, Sai Zhang

## Abstract

Plasma proteomic profiling has been widely used for biomarker discovery, disease prediction and diagnosis, and patient stratification. However, technical differences across assay platforms often result in low-to-moderate agreement, limiting study reproducibility, data integration, and model transferability. Here we present AART, a cross-platform proteomic translation framework that integrates matched-protein ridge regression with proteome-wide residual learning. We benchmarked AART spanning three independent cohorts profiled using three major platforms, including Olink, SomaScan, and mass spectrometry. Across all six translation directions, AART achieved the best performance compared with baseline methods for both overlapping and non-overlapping protein translations, with a relative improvement of 92.0% on average over direct mapping and by up to 31.6% over cpiVAE, the strongest baseline. Proteins that were accurately translated and improved by AART were enriched for extracellular, vesicle-associated, and tissue-restricted plasma biology. In downstream applications, AART improved the reproducibility of proteomic association analyses relative to direct cross-platform comparison by 75.5% for type 2 diabetes and 370.6% for Alzheimer’s disease. AART-enabled cohort integration enhanced diagnostic accuracy for amyotrophic lateral sclerosis by 92.6% compared with non-integration analysis. AART was overall one to three orders of magnitude faster than cpiVAE, facilitating biobank-scale applications. Together, these results establish AART as a fast, accurate, and scalable framework for cross-platform proteomic translation, enabling more reproducible, transferable, and integrated proteomic research.

## Introduction

Plasma proteomic profiling is rapidly transforming biomarker discovery, disease prediction and diagnosis, and patient stratification^1,2^. Proteomic signatures have been developed to diagnose and predict a wide range of complex diseases, including cardiovascular diseases^3,4^, type 2 diabetes^5^, Alzheimer’s disease^6,7^, and amyotrophic lateral sclerosis^8,9^. Proteomic aging clocks have expanded molecular hallmarks of human aging and provide quantitative indicators for disease risk and progression^10–12^. The clinical translation of these advances depends on whether protein measurements and derived signatures are interpretable across studies, cohorts, and assay platforms.

Large-scale plasma proteomic resources have been generated using different high-throughput technologies, most prominently Olink, SomaScan, and mass spectrometry (MS). Olink uses antibody-based proximity extension assays^13^, SomaScan uses modified aptamer reagents^14,15^, and MS infers protein abundance from peptide-level intensities after liquid-chromatography separation^16–18^. These technologies provide complementary views of the circulating proteome; however, direct translation of proteomic measurements across platforms remains challenging^19^. Empirical studies have consistently shown limited concordance across proteomic platforms. In the China Kadoorie Biobank (CKB), Olink and SomaScan showed only modest agreement across matched protein targets^20^, with correlations ranging from 0.26 to 0.29. Similar discordance has been reported in other Olink-SomaScan comparisons and in broader assessments involving affinity-based and MS-based platforms^21–23^. These systematic differences arise from both analyte-level biological complexity^24^ and platform-specific technical variation^20,25,26^, limiting study reproducibility, model transferability, biomarker validation, and cross-cohort integration.

Recent studies address assay heterogeneity by retraining platform-specific models or harmonizing derived phenotypes, rather than directly translating protein measurements across assays. For example, Ding *et al*. developed separate cellular aging clocks using SomaScan and Olink^27^, in which model outputs were standardized as age-gap phenotypes and extreme aging states were binarized to mitigate cross-platform variation. Oh *et al*. rebuilt organ aging models using Olink to synchronize previously developed SomaScan-based models^28,29^, along with platform-specific protein and pathway signatures. Although these approaches improve cross-platform transferability at the phenotype or model-output level, they do not resolve the underlying discordance in protein measurements.

These challenges motivate the development of computational methods for cross-platform proteomic translation. For example, cpiVAE, a joint variational autoencoder, enables bidirectional translation between Olink and SomaScan by learning a shared latent representation across the two platforms^30^. However, existing methods have largely been designed and evaluated for specific platform pairs, with a focus on proteins measured by both assays. Thus, there remains a need for a universal framework that can accurately translate protein measurements across major platforms and across the full set of measured proteins.

In this study, we developed the **<u>A</u>**nchor-**<u>A</u>**ware **<u>R</u>**esidual **<u>T</u>**ranslator (AART), a unified framework for translating protein measurements among Olink, SomaScan, and MS. AART first learns a matched-anchor ridge regression model for each target protein to capture direct cross-platform correspondence, and then models residual target-platform variation using global PCA ridge regression. Both local and global predictions are then combined via a protein-specific reliability gate, allowing AART to adaptively balance between matched-protein information and proteome-wide residual structure. Across three independent cohorts and six translation directions among Olink, SomaScan, and MS, AART outperformed all baseline methods, including Direct-1:1, KNN, WNN, and cpiVAE. We further demonstrated the utility of AART in several downstream applications, including biomarker discovery and cohort integration. AART-based translation improved analysis reproducibility and preserved disease-associated protein markers for type 2 diabetes (T2D) and Alzheimer’s disease (AD), increasing effect-size correlations from 0.17-0.53 between original platforms to 0.80-0.93 after translation. AART-enabled cohort integration improved diagnostic accuracy for amyotrophic lateral sclerosis by 92.6% compared with non-integration analysis. Importantly, AART’s simple mathematical formulation and closed-form solution enable efficient implementation, achieving an approximately 1,000-fold speedup over cpiVAE. Together, these results establish AART as a versatile, fast, and accurate framework for cross-platform proteomic translation, providing a methodological foundation for scalable and reproducible proteomic studies.

## Results

### Overview of AART

We developed AART to translate plasma proteomic profiles across multiple major platforms, including Olink, SomaScan and mass spectrometry (MS). The model was developed and benchmarked using paired measurements from the China Kadoorie Biobank^20^ (CKB) for Olink-SomaScan translation, PEX-LC^31^ for Olink-MS translation, and the Global Neurodegeneration Proteomics Consortium^32^ (GNPC) for MS-SomaScan translation, spanning all six pairwise translation directions among the three platforms (**Fig. 1a**; **Methods**). For downstream applications, GNPC data were used for proteomic association analyses, whereas UK Biobank Olink data^1^ were used for cohort integration. Protein-level and sample-level quality control (QC) procedures were conducted for all datasets before analysis (**Methods**).

**Fig. 1.**
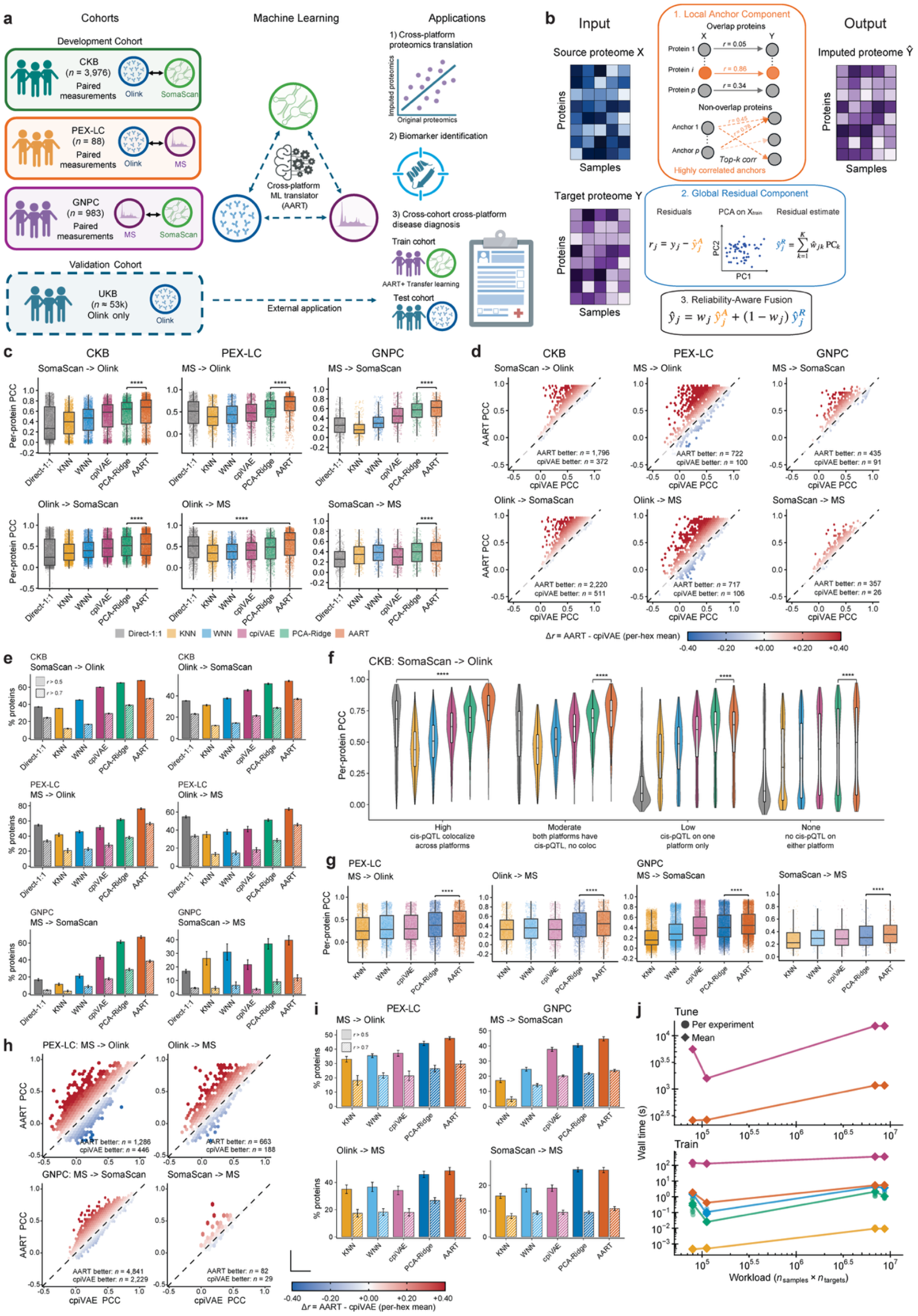
Development and benchmarking of AART for cross-platform proteomic translation. **a**, Study design. Three paired-assay cohorts were used for method development and benchmarking: CKB (*n* = 3,976) for Olink-SomaScan translation, PEX-LC (*n* = 88) for Olink-MS translation, and GNPC (*n* = 983) for MS-SomaScan translation. UK Biobank Olink data were used as an external application cohort. **b**, Schematic of AART. A source-platform proteome is translated into a target-platform proteome by combining a local anchor component, a global residual component, and a reliability-aware fusion. For overlapping proteins, the local component uses matched source-platform proteins as anchors. For non-overlapping proteins, top-*k* highly correlated source proteins are selected as anchors. The global component models target-platform residual structure using principal components of the source proteome, and a per-protein reliability gate finally integrates both local and global predictions. **c**, Cross-platform proteomic translation performance for overlapping proteins. Box plots show five-fold cross-validated per-protein Pearson’s *r* across held-out test samples between translated and observed target-platform protein expression levels. Performance was evaluated for Direct-1:1, KNN, WNN, cpiVAE, PCA-Ridge, and AART across six cohort-by-direction translation tasks. The box plot center line, limits, and whiskers represent the median, quartiles, and 1.5x interquartile range (IQR), respectively. ****, *P* < 1e-4 by one-sided paired Wilcoxon signed-rank test. All comparisons were performed between AART and the best baseline method. PCC, Pearson correlation coefficient. **d**, Protein-level comparison between AART and cpiVAE for overlapping proteins. Hexbin scatter plots compare five-fold mean values of per-protein Pearson correlations between AART and cpiVAE. **e**, Performance stratification by different correlation thresholds. Bar plots show the percentage of overlapping proteins with five-fold mean PCC values greater than 0.5 or 0.7 across methods and translation directions. The bar plot represents the mean percentage across folds, and the error bar denotes the standard error. **f**, Translation performance stratified by genetic concordance. Violin plots show per-protein PCCs for CKB SomaScan->Olink translation, grouped by cross-platform cis-pQTL colocalization tiers. The box plot center line, limits, and whiskers represent the median, quartiles, and 1.5x IQR, respectively. ****, *P* < 1e-4 by one-sided paired Wilcoxon signed-rank test. All comparisons were performed between AART and the best baseline method. **g**, Cross-platform proteomic translation performance for non-overlapping proteins. Box plots show five-fold cross-validated per-protein Pearson’s *r* across held-out test samples for GNPC (MS-SomaScan) and PEX-LC (Olink-MS) proteins without direct source-platform counterparts. Performance was evaluated for KNN, WNN, cpiVAE, PCA-Ridge, and AART. The box plot center line, limits, and whiskers represent the median, quartiles, and 1.5x IQR, respectively. ****, *P* < 1e-4 by one-sided paired Wilcoxon signed-rank test. All comparisons were performed between AART and the best baseline method. **h**, Protein-level comparison between AART and cpiVAE for non-overlapping proteins. Hexbin scatter plots show per-protein PCCs of AART versus cpiVAE for non-overlapping proteins. **i**, Performance stratification by different correlation thresholds for non-overlapping protein translation. Bar plots show the percentage of non-overlapping proteins with five-fold mean PCC values greater than 0.5 or 0.7 across methods and translation directions. The bar plot represents the mean percentage across folds, and the error bar denotes the standard error. **j**, Runtime comparison. Wall time for model hyperparameter tuning and model training is plotted against workload, defined as the product of sample size and number of proteins. Both axes are shown on a log scale. Runtime was evaluated using the same benchmark tasks across methods where applicable.

AART consists of three components: a local anchor component, proteome-wide residual learning, and protein-specific reliability-aware fusion (**Fig. 1b**; **Methods**). For overlapping target proteins measured by both assays, the local component uses identifier-matched source features as the anchor. For non-overlapping target proteins measured by only one assay, it instead uses the top correlated source features (**Methods**). To capture target-platform variation not explained by the local anchor, AART further introduces a PCA ridge model to predict the residual. The local and residual predictions are then combined through a protein-specific reliability gate. For fair comparison, all translation methods were evaluated using identical five-fold cross-validation splits, with the per-protein Pearson correlation coefficient (PCC) computed across held-out test samples as the primary performance metric.

### AART improves translation of overlapping proteins

We first evaluated AART’s performance for translating overlapping proteins, benchmarked against multiple baseline methods (**Methods**), including the direct translation (Direct-1:1), *k*-nearest neighbors (KNN), weighted-nearest neighbors^33^ (WNN), and cpiVAE^30^. We also implemented a single PCA-Ridge component of AART as an additional baseline.

Across all six translation tasks, AART outperformed all baseline methods evaluated by per-protein PCC (*P* < 2.5e-4, one-sided paired Wilcoxon signed-rank test; **Fig. 1c**). The median PCCs of AART were 0.678 and 0.554 for the two CKB directions, 0.749 and 0.706 for the two PEX-LC directions, and 0.617 and 0.406 for the two GNPC directions. Notably, PCA-Ridge, a component of AART, was the second-best-performing method for all translation tasks except Olink->MS. Relative to the best baseline method in each task, AART improved median PCC by 0.3% to 31.6%, with the largest gains observed for MS-Olink translation in PEX-LC.

Protein-wise comparisons showed that AART’s performance gains were broadly distributed across proteins. Across the six translation tasks, AART achieved higher Pearson correlations than cpiVAE for approximately 80-93% of overlapping proteins (**Fig. 1d**). Moreover, AART increased the number of proteins reaching predefined performance thresholds (**Fig. 1e**). For example, the proportions of proteins with five-fold mean *r* > 0.5 by AART ranged from 39.6% to 78.3% across the six translations, whereas the proportions with *r* > 0.7 ranged from 10.9% to 59.9%. AART achieved the highest proportion at both thresholds in every task.

The improvement was robust across alternative evaluation metrics. AART achieved the best performance across the six translations when evaluated by per-protein Spearman correlation, as well as by sample-level Pearson and Spearman correlations (**Extended Data Fig. 1a–c**). The translation gains achieved by AART for individual proteins (**Extended Data Fig. 2**), including established proteomic markers such as NEFL^8^ and LCN2^34^, further underscore its utility in biomarker studies.

### Genetic concordance stratifies proteomic translation

We next examined whether translation accuracy varied with the consistency of genetically regulated proteins captured across different platforms. Genetic regulation of protein expression was assessed using the cis protein quantitative trait locus (cis-pQTL) analysis^20^. Specifically, we grouped CKB SomaScan-Olink overlapping proteins into four tiers: proteins with colocalized cis-pQTLs across platforms, proteins with cis-pQTLs identified on both platforms but without colocalization, proteins with cis-pQTLs identified on only one platform, and proteins with no cis-pQTL identified on either platform (**Fig. 1f**).

Proteins with colocalized cis-pQTLs were generally well translated (SomaScan->Olink) by both direct mapping and multivariate methods, whereas the advantage of AART increased as shared genetic signals weakened. Direct-1:1 performance declined markedly for proteins with cis-pQTLs on only one platform or no cis-pQTL on either platform. In contrast, AART retained substantially higher predictive accuracy, suggesting that it captured variation beyond direct cross-platform protein correspondence. AART also achieved the highest per-sample correlations across strata defined by cis-pQTL colocalization, trans-pQTL colocalization, and linkage disequilibrium between cross-platform cis-pQTL signals (**Extended Data Fig. 3a–c**). Similar patterns were observed for Olink->SomaScan translation (**Extended Data Fig. 4**).

### AART extends translation to non-overlapping proteins

The sets of proteins profiled by different platforms differ substantially, and previous cross-platform analyses have typically focused on proteins measured by both assays^20,30^. We next sought to assess AART’s ability to translate proteins measured by only one platform. For benchmarking, we extended all baseline methods except Direct-1:1 to non-overlapping protein translation and focused on Olink-MS and SomaScan-MS translation tasks given data availability. In general, non-overlapping targets were less accurately translated than overlapping proteins. Nevertheless, AART achieved the best performance across all translation directions compared with baseline approaches (**Fig. 1g**), followed by PCA-Ridge. Consistent with the overlapping-protein analyses, AART’s gains were broadly distributed across proteins: AART achieved higher PCCs than cpiVAE for 69-78% of non-overlapping targets (**Fig. 1h**). AART also increased the proportion of proteins exceeding PCC thresholds of 0.5 and 0.7 across all four tasks (**Fig. 1i**). These results were stable across alternative evaluation metrics (**Extended Data Fig. 5**). Notably, several well-known protein markers such as IGFBP3^35^ and APOE4^36^ were accurately translated by AART despite being absent from the source platforms (**Extended Data Fig. 6**), highlighting its capacity to recover a more comprehensive proteome landscape for biomarker discovery.

### AART scales efficiently across cohort-size workloads

We further assessed the computational complexity of AART by measuring wall time for hyperparameter tuning, model training, and inference across workloads defined by the product of sample size and number of target proteins. AART was overall one to three orders of magnitude faster than cpiVAE, while maintaining runtimes close to those of simpler linear baselines (**Fig. 1j, Extended Data Fig. 7**). Its computational cost was dominated by ridge-regression fitting, principal-component projection, and estimation of per-protein reliability weights. These results support a reference-based translation strategy, in which AART is trained on moderately sized paired-platform cohorts and then applied to large-scale single-platform cohorts for proteomic translation.

### AART-translatable proteins are enriched for extracellular and tissue-restricted plasma biology

We next characterized the functional properties of proteins that were well translated by AART (**Methods**). Among overlapping proteins, well-translated proteins (*r*_AART_ ≥ 0.7) were consistently enriched for extracellular and vesicle-associated cellular component terms across six translation directions, including extracellular exosome, extracellular vesicle, and extracellular organelle (**Fig. 2a**, left), with the exception of SomaScan->MS translation, probably reflecting the lower number of well-translated proteins (*n* = 27). Interestingly, well-translated non-overlapping proteins (*r*_AART_ ≥ 0.7) exhibited a distinct enrichment pattern, with over-representation of cytoplasm, cytosol, nucleus, and cytoskeleton terms (**Fig. 2a**, right). We also examined overlapping proteins for which AART increased PCCs by at least 0.30 relative to Direct-1:1. These proteins were enriched for markers differentially expressed in specific tissues, including brain, smooth muscle, and lymphoid tissue (**Extended Data Fig. 8**). These results suggest that the translatability of circulating proteins may be related to their cellular localization and tissue-specific expression patterns, although further validation is needed.

**Fig. 2.**
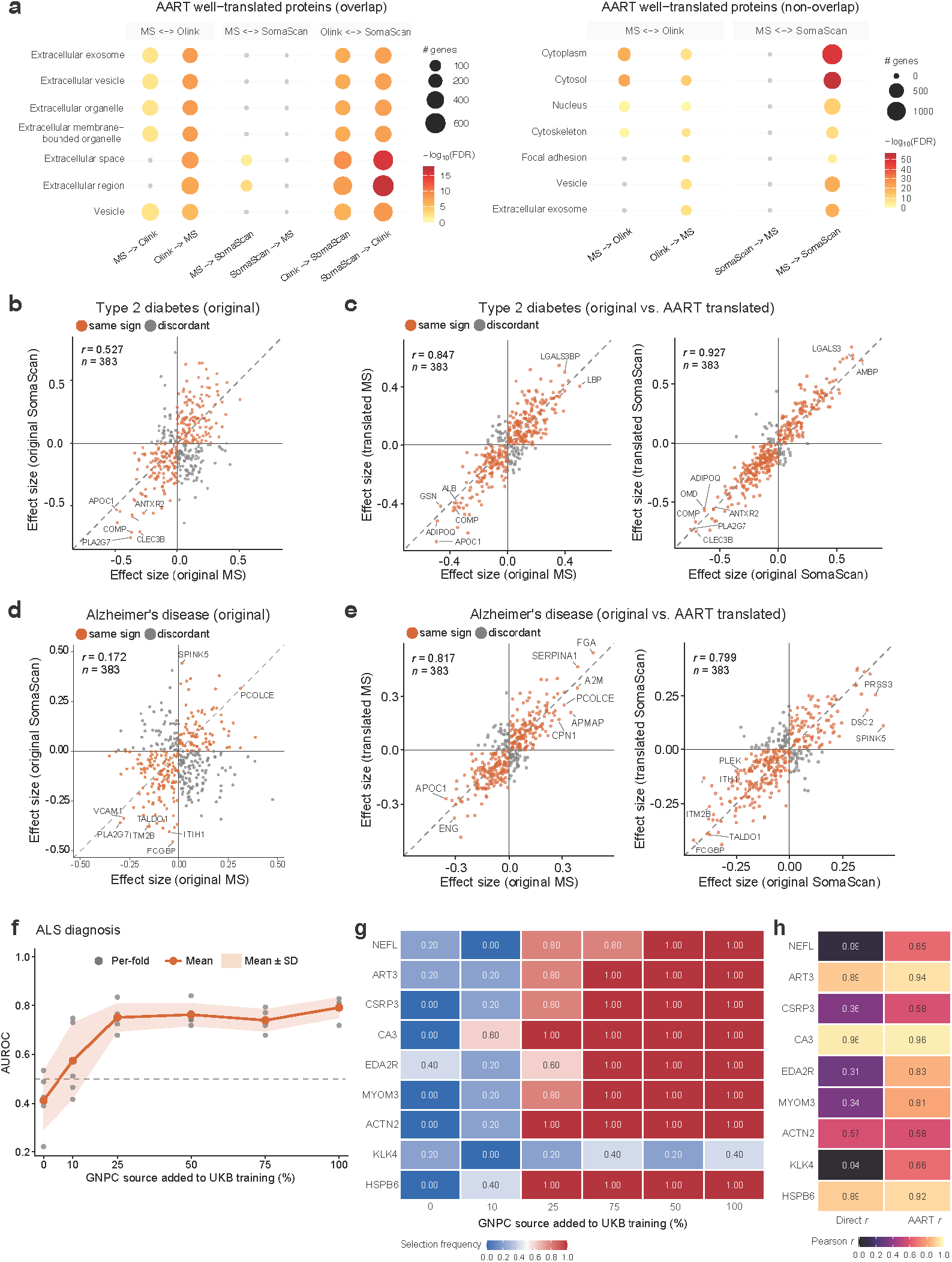
Biological characterization of AART-translated proteins and downstream applications of AART. **a**, Gene ontology (GO) enrichment of AART-translatable proteins. Bubble plots present GO enrichment analyses for well-translated overlapping (left) and non-overlapping proteins (right). Bubble color denotes −log_10_(FDR), grey bubbles denote non-significance, and bubble size indicates number of genes. MS, mass spectrometry. **b-c**, Type 2 diabetes-associated protein effects estimated before (**b**) and after AART translation (**c**). Scatter plots compare effect estimates derived from different settings. **d-e**, Alzheimer’s disease-associated protein effects estimated before (**d**) and after AART translation (**e**). Scatter plots compare effect estimates derived from different settings. In **b-e**, analyses were performed using the SomaScan-MS paired GNPC subset (*n* = 366) and 383 overlapping proteins. Each dot represents one protein; dashed lines indicate perfect match; orange dots denote concordant effect directions and grey dots denote discordant directions. Representative proteins with large absolute effects, including several proteins previously implicated in metabolic, inflammatory, vascular, or neurodegenerative diseases, are highlighted. **f**, ALS classification performance in different data augmentation settings. Grey dots denote per-fold AUROC; the orange line represents the mean and the shaded band shows the standard deviation (SD). ALS, amyotrophic lateral sclerosis; AUROC, area under the receiver operating characteristic. **g**, ALS-associated plasma proteins selected by abundance analysis. The heatmap shows selection frequencies of proteins across GNPC-augmentation fractions. **h**, Translation correlations for ALS-associated plasma proteins. The heatmap shows Pearson correlations of CKB SomaScan->Olink translation based on Direct-1:1 and AART.

### AART enhances the reproducibility of proteomic association analysis

Limited concordance among proteomic profiling assays remains a major barrier to the reproducibility of proteomic association studies^19,37–39^. We therefore examined whether AART-translated proteomes could improve the cross-platform consistency of protein-phenotype association analyses. Specifically, we conducted plasma proteomic association analyses for type 2 diabetes and Alzheimer’s disease (**Methods**), comparing results from observed proteomic measurements with those from AART-translated profiles. Among 383 overlapping UniProt-level proteins in the paired GNPC dataset, type 2 diabetes effect estimates derived from the original MS and SomaScan measurements showed only moderate agreement (Pearson’s *r* = 0.527; **Fig. 2b**). By contrast, association coefficients estimated from AART-translated profiles closely matched those from the corresponding observed target platforms, with correlations of 0.847 for MS and 0.927 for SomaScan (**Fig. 2c**), representing an increase in cross-platform reproducibility of up to 1.76-fold. Proteins significantly associated with type 2 diabetes in observed and translated analyses for the same platform also exhibited substantial overlap (**Extended Data Fig. 9a**).

The improvement was particularly pronounced for Alzheimer’s disease. Effect estimates derived directly from the two original platforms showed weak cross-platform agreement (Pearson’s *r* = 0.172; **Fig. 2d**), whereas correlations between observed and AART-translated effects reached 0.817 for MS and 0.799 for SomaScan (**Fig. 2e**), representing an increase in reproducibility of up to 4.75-fold. Significant Alzheimer’s disease-associated proteins also markedly overlapped between observed and translated analyses (**Extended Data Fig. 9b**).

We further extended these analyses to non-overlapping protein sets. As expected, disease-effect concordance between observed and translated proteins was lower than that for overlapping proteins, consistent with the greater difficulty of non-overlapping protein translation. Nevertheless, observed and translated protein effects remained positively correlated for type 2 diabetes, with correlations of 0.552 across 111 MS non-overlapping proteins and 0.587 across 6,042 SomaScan non-overlapping features (**Extended Data Fig. 10a**). For Alzheimer’s disease, the corresponding effect correlations were 0.279 for MS non-overlapping proteins and 0.510 for SomaScan non-overlapping features (**Extended Data Fig. 10b**). UpSet analyses further demonstrated overlap among nominally significant non-overlapping proteins identified from observed and translated profiles (**Extended Data Fig. 10c,d**). Together, these results underscore that AART enables reliable cross-platform proteomic association analysis, thereby enhancing the reproducibility and scope of biomarker discovery.

### AART enables cross-cohort integration

Finally, we evaluated the utility of AART for integrating proteomic cohorts profiled by different assay platforms. Specifically, a SomaScan->Olink AART translator trained on CKB was applied to the GNPC ALS cohort, comprising 355 samples including 245 ALS patients profiled by SomaScan. We then incrementally added translated GNPC profiles to the UK Biobank Olink training data (*n* = 19 ALS patients), and assessed ALS case-control classification (i.e., ALS diagnosis) based on plasma proteomic profiles (**Methods**). Classifiers were trained using augmented datasets containing increasing fractions of translated GNPC samples, and performance was evaluated on held-out UK Biobank samples (*n* = ∼5 ALS patients).

Notably, mean held-out AUROC (area under the receiver operating characteristic) increased from 0.41 using UK Biobank data alone to approximately 0.75 after incorporating 25% of the translated GNPC samples, and further to 0.79 when the full translated GNPC cohort was included (**Fig. 2f**). The performance gain was concentrated in the first 25% of source augmentation, with AUROC remaining relatively stable thereafter.

Source-cohort augmentation also improved the statistical power of ALS-associated protein marker discovery. Chia *et al*. previously identified a 17-protein Olink-based candidate biomarker panel for ALS classification^8^. 12 proteins from this panel were presented in our SomaScan-Olink overlapping proteins. Nine of them including NEFL, ART3, CSRP3, CA3, EDA2R, MYOM3, ACTN2, KLK4 and HSPB6 were prioritized in most or all cross-validation folds after incorporating translated GNPC samples (**Fig. 2g**), whereas their selection was less consistent when using UK Biobank data alone. This enhanced biomarker recovery was accompanied by higher SomaScan->Olink translation correlations for proteins with weak direct cross-platform correspondence (**Fig. 2h**), including NEFL (*r* = 0.09 to *r* = 0.65), EDA2R (*r* = 0.31 to *r* = 0.83), MYOM3 (*r* = 0.34 to *r* = 0.81) and KLK4 (*r* = 0.04 to *r* = 0.66). Proteins with strong direct correspondence, including ART3 (*r* = 0.89), CA3 (*r* = 0.96), and HSPB6 (*r* = 0.89), retained high correlations after AART translation. Together, these results demonstrate that AART-enabled cohort integration improves biomarker selection stability and disease classification accuracy, thereby increasing the statistical power of proteomic studies for rare diseases.

## Discussion

We present AART, a fast, accurate, and scalable framework for cross-platform proteomic translation. Large-scale plasma proteomic datasets are increasingly available from Olink, SomaScan, and mass spectrometry across population biobanks and disease-specific cohorts. These technologies measure partially overlapping but heterogeneous protein feature spaces, in which direct cross-platform correspondence can be confounded by batch effects and platform-specific technical biases. The impact of this heterogeneity extends beyond protein abundance measurements: previous comparative studies have reported limited replication of genetic and phenotypic associations across proteomic platforms^40,41^, and our GNPC analyses similarly revealed low-to-moderate agreement between MS-and SomaScan-derived protein-disease associations. AART addresses this challenge by learning proteome-wide mappings from paired-platform reference cohorts and translating single-platform measurements into the target-platform representation. Across three independent cohorts and six translation directions, AART consistently outperformed a broad range of baseline methods, achieving state-of-the-art translation performance. This superiority extended to both overlapping and platform-specific non-overlapping proteins, substantially expanding the scope of cross-platform proteomic integration.

A defining feature of AART is its reliability-aware design. On one hand, direct one-to-one mapping is simple and interpretable, but becomes unreliable when nominally matched proteins show weak cross-platform correspondence or when differences in reagents, peptides, or detection windows capture non-identical molecular signals. On the other hand, models built on global proteome-wide structure, such as cpiVAE, can leverage multivariate covariance but may underuse strong protein-specific anchors when they are available. AART reconciles these two strategies. For each target protein, the matched-anchor ridge component captures direct correspondence between the source and target platforms, whereas the residual PCA-ridge component models target-platform variation that remains unexplained by the matched anchors. A protein-specific reliability gate then integrates the local anchor prediction with the global residual correction. This design allows AART to preserve interpretable protein-specific signals when direct correspondence is reliable, while borrowing information from broader proteome-wide covariance when matched-anchor information is insufficient. The cis-pQTL analyses support this interpretation: proteins with strong cross-platform genetic concordance were generally better translated, whereas AART provided additional gains for proteins with weaker direct genetic or matched-anchor signals. Thus, AART should not be interpreted as assuming that all platform readouts are equivalent. Rather, it offers a reliability-aware framework for estimating plasma protein variation that is statistically and biologically transferable across assay platforms.

Non-overlapping protein translation extends AART beyond the harmonization of proteins shared by two platforms. Platform-specific proteins can have significant biological and clinical value^42^, complementing overlapping proteins to offer a more complete view of the proteomic landscape. In both PEX-LC and GNPC, many target features lacked a UniProt-matched source-platform measurements after quality controls and feature matching. These targets were less accurately translated than overlapping proteins, as no direct matched anchor was available. To address this, AART used source proteins selected by their correlations with each target as similarity-based pseudo-anchors, together with target-space structure and proteome-wide residual learning. This strategy improved non-overlapping protein translation in both directions between MS and Olink in PEX-LC and between MS and SomaScan in GNPC, outperforming all baseline methods.

Proteins well translated by AART exhibited coherent biological functions. Accurately translated overlapping proteins were enriched for extracellular and vesicle-associated compartments, and proteins with the largest improvements over Direct-1:1 were enriched for tissue-specific proteins. Well-translated non-overlapping targets were enriched for intracellular compartments and coordinated cellular modules. Proteins within these compartments often participate in shared metabolic, structural, trafficking or secretory processes, and may therefore present coordinated abundance across individuals owing to common tissue origin, cellular turnover or pathway activity^43^. These findings suggest that AART may recover biologically structured variation embedded in the circulating proteome, while highlighting the need for further experimental validation of the underlying regulatory mechanisms.

AART is computationally efficient and well suited for biobank-scale applications. Unlike deep learning approaches that require extensive hyperparameter tuning and iterative optimization, AART relies on ridge regression, principal-component projection, and analytically solvable reliability gates. After the anchor models, residual models, and reliability weights are learned, translation of new samples reduces to lightweight matrix operations. This design made AART one to three orders of magnitude faster than cpiVAE across the evaluated runtime settings, while keeping its computational cost close to that of simple linear baselines.

Two downstream applications highlight the utility of AART. Limited reproducibility of proteomic association analyses across platforms remains a major challenge for proteomic research. We demonstrated that AART-based protein translation can mitigate this issue and substantially enhanced the replication of biomarker discoveries. In GNPC, protein effect estimates derived directly from the original MS and SomaScan measurements showed only low-to-moderate agreement for type 2 diabetes and Alzheimer’s disease. This agreement was markedly improved by up to 4.75-fold when association analyses were performed using AART-translated proteins. In a proteomics-based ALS diagnosis task, AART-enabled cohort integration improved diagnostic accuracy by 1.9-fold, from 0.41 to 0.79. This data augmentation strategy also boosted the stability and power of biomarker discovery, reinforcing the advantage of cross-platform cohort integration. Although ALS served as the demonstration case, the same strategy can be broadly applied to clinical proteomic studies in which limited sample size, low disease prevalence, or platform difference restrict cohort-specific analyses.

Deep learning and foundation models are transforming biomedical research. However, recent studies have shown that complex models do not always outperform simpler linear models across a range of tasks, including personal genotype-gene expression prediction^44,45^, single-cell analysis^46^, and perturbation-effect prediction^47^. One possible explanation for this gap between expected and observed performance is that the amount of effective training data currently available for these tasks, such as perturbation measurements across contexts and individual-level single-cell profiles, remains insufficient to train large models to their full potential or to reach favorable scaling regimes. In our setting, proteomic cohort sample sizes are still relatively modest compared with those available in other domains. It is therefore not unexpected that AART, a structured linear model designed to exploit cross-platform proteomic data structure, yields better generalizability and outperforms deep learning approaches such as cpiVAE.

Our framework has several limitations. First, the current implementation of AART supports supervised translation only for proteins that are measured, or can be matched, across platforms in paired reference data. It therefore does not perform *de novo* prediction of proteins absent from the target assay, nor does it expand coverage to features for which no cross-platform training data is available. Second, AART estimates target-platform variation that is statistically recoverable from the source platform. Accordingly, AART-translated profiles are best viewed as platform-aligned estimates, rather than substitutes for direct assay measurements in settings requiring analyte-specific quantification.

In conclusion, AART establishes a foundational framework for translating plasma proteomic measurements across assay platforms. By connecting single-platform cohorts, AART offers a practical path toward more reproducible and interoperable proteomic research. We provide AART as an open-source implementation with tutorials and reproducible workflows for model training, benchmarking, and application to new cohorts.

## Data availability

The CKB and PEX-LC proteomic datasets can be accessed through their original publications^20,31^. The GNPC V1 harmonized dataset (HDS) is available upon request, with request instructions provided at https://www.neuroproteome.org/harmonized-data-set-hds. UK Biobank proteomic data are available to registered researchers through the UK Biobank Access Management System (https://www.ukbiobank.ac.uk), subject to approved access; this study was conducted under UK Biobank application number 101835.

## Code availability

The source code, tutorials, and reproducible workflows of AART are available at https://github.com/saizhanglab/AART.

## Acknowledgments

This study was supported by the NIGMS (R35GM157219 to S.Z.), the Motor Neurone Disease Association (to S.Z.), the Packard Center for ALS Research (to S.Z.), and the Target ALS Foundation (to S.Z.).

## Author contributions

Y.C. and S.Z. conceived and designed the study. Y.C. developed AART and performed the data analyses. Y.C. and S.Z. interpreted the results. S.Z. supervised the project. Y.C. and S.Z. wrote the manuscript.

## Methods

### Machine learning formulation for cross-platform proteomic translation

To make proteomic measurements interoperable across assay technologies, we formulated cross-platform translation as a supervised-learning problem. For each individual *i* in a paired cohort, the source-platform proteome is represented as **x**_*i*_ ∈ ℝ^*pS*^ and the target-platform proteome **x**_*i*_ ∈ ℝ^*pS*^, where *p*_*S*_ and *p*_*T*_ denote the numbers of proteins measured on the source and target assays, respectively. Given a training set of *n* paired samples 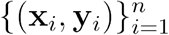, we learn a parameterized translator:

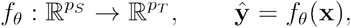

by minimizing the mean squared reconstruction error between predicted and observed target-platform values,

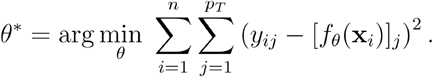

The trained translator 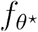 is then applied to unseen individuals profiled on the source platform, producing a translated target proteome 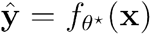 for use in downstream analyses.

### The AART model

AART is an anchor-augmented residual translator trained on paired cross-platform reference dataset. For a given translation direction, let 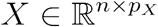 denote the standardized source-platform matrix and 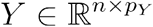 the standardized target-platform matrix measured in the same *n* samples. AART learns a translation function *f* : *X* → *Y* by defining, for each target protein, the most correlated anchor in the source space and then modeling the residual target-platform variation using regularized regression.

For the target feature *j* in sample *i*, AART’s prediction is given by

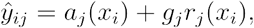

where *a*_*j*_(*x*_*i*_) is the anchor prediction, *r*_*j*_(*x*_*i*_) is the residual correction learned from source-platform proteome-wide structure, and *g*_*i*_ is a target-specific reliability gate controlling the contribution of the residual component. Each component is defined separately for overlapping targets with a direct source-platform counterpart and non-overlapping targets without such a counterpart.

#### Step 1: Anchor-based prediction

For each overlapping target feature *j* ∈ *O*, let *s*(*j*) denote the identifier-matched source feature used as its direct anchor. Features are matched using shared UniProt identifiers or platform-specific mapping. A target-specific ridge regression is fitted across training samples *T*:

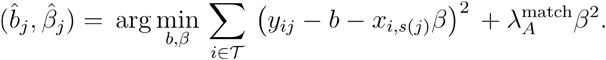

The resulting anchor prediction is then given by

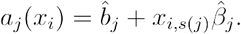

This anchor provides a direct, target-specific estimate based on the source-platform measurement.

For the non-overlapping target feature *j* ∈ *N*, since no identifier-matched source feature is available, AART constructs an ensemble anchor using two complementary strategies: (1) a target-space PCA prediction that captures shared low-rank structure across target features and (2) a correlation-based prediction that captures target-specific relationships with source features. Specifically, AART first translated overlapping targets and appends their predicted values to the original source matrix:

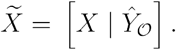

Here, the translated overlapping targets are used as auxiliary features for predicting non-overlapping targets. Next, principal component analyses are performed for both the augmented source matrix and the non-overlapping target matrix:

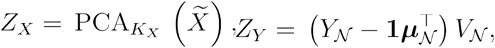

in which ***μ***_*N*_ is the target feature mean vector and *V*_*N*_ represents the retained target-space principal-component loadings. A multivariable ridge regression is fitted from the augmented source PC feature *Z*_*X*_ to the target PC feature *Z*_*Y*_, and the prediction is obtained in the original target-feature space:

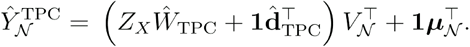

We further expand the anchor estimation by combining the correlation-based protein predictions. For each non-overlapping target *j*, the original source features are ranked by their absolute Pearson correlations with the target across the training samples:

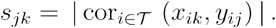

The *K*_*C*_ highest-ranking source features define a target-specific pseudo-anchor set:

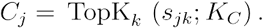

A target-specific ridge regression is fitted using these selected source features:

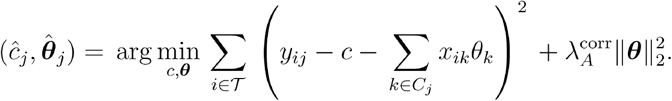

The correlation-anchor prediction is given by

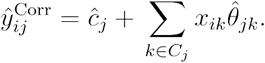

Finally, the PCA-based and correlation-anchor predictions are combined to yield the non-overlapping protein prediction:

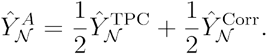

The resulting overlapping and non-overlapping protein predictions are subsequently passed to the residual-learning component.

#### Step 2: Proteome-wide residual learning

The residual component models target-platform variation that is not captured by the anchor prediction. For both overlapping and non-overlapping targets, AART defines the residual matrix as the difference between the observed target measurements and anchor-based predictions. It then fits a multivariable PCA-ridge model to predict these residuals. The two settings differ in the source representation adopted by the residual model.

For the overlapping target set *O*, 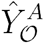 let denote the direct matched-anchor predictions. The anchor residuals are defined as

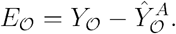

The original source matrix is projected onto its first 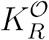 principal components:

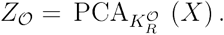

A multivariable ridge regression is then fitted to map the source principal components to the anchor residuals:

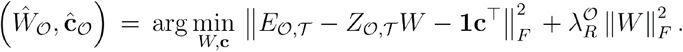

The resulting residual predictions are given by

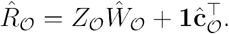

For the non-overlapping target set *N*, 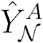 let denote the ensemble anchor predictions obtained from the PCA-based and correlation-anchor components. The corresponding residuals are therefore given by

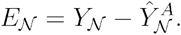

Because no directly matched source feature is available for these targets, the residual regression model uses the augmented source matrix constructed in Step 1:

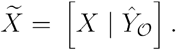

A separate principal-component representation is estimated from the augmented source matrix:

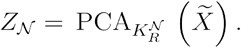

Similarly as the design for overlapping protein prediction, a multivariable ridge regression is then fitted from the augmented source components to the non-overlapping anchor residuals:

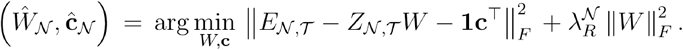

The residual predictions for non-overlapping targets are given by

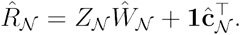

For overlapping targets, the residual component leverages the global structure in the original source proteome space. For non-overlapping targets, it additionally uses the translated overlapping targets as auxiliary features. In both settings, the residual model captures proteome-wide co-abundance structure and target-platform variation not represented by the anchor component.

#### Step 3: Reliability-aware gating

For both overlapping and non-overlapping targets, anchor and residual models are fitted in the training set *T*, whereas the target-specific gate is estimated in a separate validation set *V*.

For a given target protein *j*, the gate can be obtained analytically as

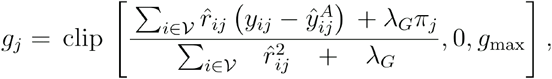

where 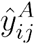 is the validation-set anchor-based prediction, 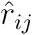 is the residual prediction, *π*_*j*_ is a branch-specific prior residual weight, *λ*_*j*_ controls shrinkage toward this prior, and *g*_max_ defines the maximum permitted residual correction. For overlapping targets, the prior is determined by the reliability of the directly matched anchor. Specifically, let

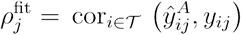

denote the Pearson correlation between the anchor-based prediction and observed target values in the training set. The prior residual weight is given by

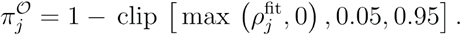

Thus, a strongly predictive direct anchor receives a small prior residual contribution, whereas a weakly or negatively correlated anchor receives a large residual contribution. The overlapping-target gate is constrained to

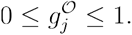

The final overlapping-target prediction is given by

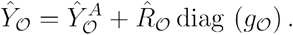

Because non-overlapping targets lack a direct identifier-matched anchor, their anchor reliability cannot be defined using direct cross-platform correspondence. The non-overlapping branch therefore uses a common prior residual weight:

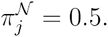

The gate is estimated using the same closed-form equation, and the final non-overlapping-target prediction is given by

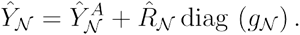

For either branch, a gate value of zero gives the anchor-only prediction, and a gate value of one adds the full residual prediction.

All hyperparameters, data standardization, identifier maps, similarity scores, selected anchor features, PCA loadings, ridge coefficients, and gates were estimated and determined within the training data for each cross-validation fold. At the inference time, a new source sample was standardized based on the training data, passed through the AART model, and then translated to the target-platform scale. No optimization was performed during inference.

### Implementation details for AART and baseline methods

Identifier-based source-target mappings were defined based on the platform annotations before cross-validation and were held fixed across folds. For each cohort and translation direction, we used the five-fold cross-validation. All data-dependent preprocessing, model selection, and model training were conducted without access to the held-out test datasets. The validation set was used for hyperparameter selection and estimation of the target-specific reliability gates. The held-out test set was used only for final performance evaluation. At inference, a new source-platform sample was standardized using statistics estimated from the reference training data.

For the overlapping-target branch, four hyperparameters for AART were selected by grid search: the matched-anchor ridge penalty, the residual ridge penalty, the number of source principal components, and the gate regularization parameter. The search spaces were 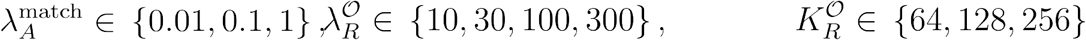, and 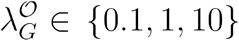. All combinations were evaluated, yielding 3 × 4 × 3 × 3 = 108 candidate configurations per outer fold. For each configuration, the anchor-based and residual models were fitted in the training set, and the reliability gates were estimated in the validation set. The configuration with the highest median per-target Pearson correlation in the validation set was selected. In analyses including non-overlapping targets, the overlapping-target branch was first fitted and selected using the procedure described above. Parameters specific to the non-overlapping branch were fixed across cohorts and translation directions.

For direct 1:1 mapping, when a target feature had more than one candidate source feature, such as an Olink protein mapping to multiple SomaScan aptamers, the candidate with the highest Pearson correlation in the validation subset was selected as the target prediction. PCA-Ridge projected source features into 256 principal components and fitted Ridge regression (alpha = 300). KNN used *k* = 10 distance-weighted neighbors, and WNN followed the Seurat-style weighted nearest-neighbor framework with 50 PCA components per modality and Gaussian kernel weighting over 50 neighbors. For cpiVAE, we performed Bayesian hyperparameter optimization using Optuna (50 trials), searching over latent dimension (*n* = 16, 32, 64, 128, 256), encoder and decoder architectures (1-3 layers of 64, 128, 256, 512, or 1024} units), learning rate (1e-5 to 1e-2), dropout rate (0.1 to 0.5), activation function (ReLU, Leaky ReLU, GELU, or Swish), batch size (*n* = 32, 64, 128, 256), loss weights for reconstruction, KL divergence, cross-reconstruction, and latent alignment, and alignment type (MSE, KL divergence, or MMD). Trials were evaluated by mean bidirectional cross-reconstruction correlation on the held-out validation set, with early stopping (patience = 10). The best configuration was then applied to train all five folds for 200 epochs with early stopping on validation loss.

### Benchmarking methods, cross-validation, and evaluation metrics

We benchmarked AART against five baseline methods spanning direct matching, local neighbor transfer, global linear mapping, and a deep learning model. Direct-1:1 used a single best matched source feature for each target protein: the best matched SomaScan aptamers for CKB Olink targets, and the shared UniProt features for MS-Olink and MS-SomaScan translation. *K*-nearest neighbors (KNN) algorithm used distance-weighted *k*-nearest-neighbor regression with *k* = 10 in the standardized source-feature space. Weighted nearest neighbors (WNN) algorithm followed the Seurat-style weighted-nearest-neighbor idea, using 50-dimensional platform-specific PC embeddings, cross-modality nearest-neighbor graphs, Jaccard-overlap weights, and a Gaussian-kernel regression step to predict target abundances from source profiles. PCA-Ridge used the top 256 source-platform PCs in a multivariable ridge regression to predict all target proteins jointly; this was the global residual component of AART without matched-anchor models or reliability gates. cpiVAE was included as a deep learning baseline^30^. Following the cpiVAE framework, dual platform-specific encoders mapped two platform profiles into a shared latent space, and platform-specific decoders reconstructed the target-platform proteome. In our benchmarks, cpiVAE was trained separately within each training fold and evaluated on the same held-out fold as used by all other methods. We adapted all methods including cpiVAE to non-overlapping protein translation, except Direct-1:1.

For each cohort and translation direction, all methods were trained and evaluated using the same five-fold cross-validation splits. The primary metric was the per-target Pearson correlation coefficient (PCC) and Spearman correlation coefficients (SCC) between translated and observed target-platform measurements. For target feature *j* in the held-out fold *k*, PCC was calculated as

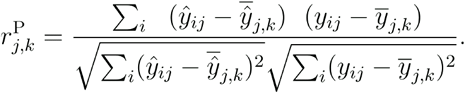

Per-target SCCs were calculated between the within-fold ranks of translated and observed values:

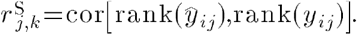

For each translation direction, per-target performance was summarized using the median, mean, and interquartile range of the cross-validated correlations. Sample-level Pearson and Spearman correlations were calculated analogously across target features within each held-out sample. Protein proportion above correlation thresholds was calculated separately in each fold as the percentage of targets with PCC greater than 0.5 or 0.7 and then averaged across the five folds. Method-specific improvement was summarized as

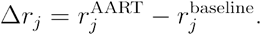

One-sided paired Wilcoxon signed-rank tests were used to compare AART with baselines across target features.

### Plasma proteomic profiling platforms

We analyzed three major high-throughput plasma proteomic profiling assays: Olink, SomaScan, and mass spectrometry. Olink Explore 3072 is an affinity-based proximity extension assay in which paired antibodies bind predefined protein targets and generate DNA reporters quantified by next-generation sequencing. Olink measurements are reported as normalized protein expression (NPX) values on a log2 scale after assay-level quality control and normalization. SomaScan is an aptamer-based assay that uses slow off-rate modified aptamers (SOMAmers) to bind proteins in solution. SomaScan v4.1 measures approximately 7,000 protein targets and reports relative fluorescence units after plate-level calibration and normalization; SomaScan values were log-transformed upstream in the source quality-control pipelines before AART modeling^20,48^. MS measurements were analyzed as protein-level abundance matrices derived from peptide evidence after study-specific quality control, missingness filtering and protein annotation.

Cross-platform features were matched by UniProt identifiers provided by the original studies. For Olink-SomaScan translation in CKB, Olink reagents and SOMAmers were mapped to UniProt IDs as described in the original platform-comparison study^20^. If multiple SOMAmers mapped to the same Olink protein target, AART retained all mapped SOMAmers as candidate anchors. For MS-Olink and MS-SomaScan translation, matched anchors were defined using shared UniProt identifiers between the protein-level MS matrix and the affinity-platform matrix. Features with matched source-platform counterparts were treated as overlapping targets, whereas target proteins without a direct counterpart on the source platform were treated as non-overlapping targets and evaluated separately. For the Direct-1:1 baseline, when multiple candidate source features mapped to the same target protein, the source feature with the highest Pearson correlation in the validation split within each training fold was selected.

Because platforms differ in molecular recognition chemistry, calibration, dynamic range, missingness structure, and molecular species captured, matched UniProt identifiers were not assumed to represent interchangeable measurements. Instead, AART treated matched features as candidate anchors and learned their predictive contribution from paired-platform reference samples.

### Study cohorts

We assembled three paired plasma proteomic datasets spanning pairwise combinations of three major assay technologies: the China Kadoorie Biobank (CKB) for Olink-SomaScan translation^20^ (*n* = 3,976; 2,168 Olink proteins and 2,731 SomaScan aptamers); the PEX-LC lung-cancer cohort for Olink-MS translation^31^ (*n* = 88; 2,780 Olink proteins and 1747 MS proteins), and the Global Neurodegeneration Proteomics Consortium (GNPC) for SomaScan-MS translation^32^ (*n* = 366; 494 MS proteins and 7,596 SomaScan aptamers). Each paired cohort supports bidirectional prediction, yielding six cross-platform translation tasks in total (**Fig. 1a**). For downstream cohort integration analysis, we used the UK Biobank Pharma Proteomics Project^1^ (UKB-PPP) Olink dataset (*n* ≈ 53k) as an external cohort.

### China Kadoorie Biobank

The China Kadoorie Biobank (CKB) is a prospective population study of more than 512k adults recruited from ten geographically diverse regions of China. The Olink-SomaScan platform comparison study profiled baseline plasma from an ischemic heart disease subcohort of 3,976 participants using both Olink Explore 3072 and SomaScan v4.1^20^. The original study reported 2,923 unique Olink reagents and 7,301 UniProt-annotated SOMAmer reagents, yielding 2,749 Olink-SomaScan reagent pairs mapped to 2,168 UniProt IDs. When multiple SOMAmers mapped to the same Olink target, all candidate SOMAmer anchors were retained for AART. CKB was used to evaluate SomaScan->Olink and Olink->SomaScan translation and to train the SomaScan->Olink translator used for the downstream cohort-integration analysis.

### PEX-LC

PEX-LC is a retrospectively selected plasma cohort of patients referred to Karolinska University Hospital for investigation of suspected lung cancer^31^. The original study analyzed 114 pre-diagnostic plasma samples using HiRIEF LC–MS/MS, with six samples run in duplicate, and profiled a subset of 88 samples using Olink Explore 3072. We used these 88 shared samples for Olink-MS translation. In the published PEX-LC comparison study, MS detected 2,578 unique proteins and Olink quantified 2,913 proteins after exclusion of Olink assays below the limit of detection across samples. The two platforms shared 1,129 UniProt IDs.

### Global Neurodegeneration Proteomics Consortium

The Global Neurodegeneration Proteomics Consortium (GNPC) is a harmonized neurodegeneration proteomics resource containing biofluid proteomic profiles from clinically healthy individuals and participants with neurodegenerative diseases^48^. SomaScan was the primary profiling technology in GNPC, with the v4.1 assay measuring approximately 7,000 aptamers per biosample. A total of 983 GNPC plasma samples had matched SomaScan and MS proteomic measurements. After dropping samples with a high missing rate for either platform, 366 samples are kept in the final analysis. In this matched subset, 383 MS proteins and 526 SomaScan aptamers had a shared UniProt ID. The remaining 111 MS proteins and 7,070 SomaScan aptamers had no match and were treated as non-overlapping features. GNPC was used to evaluate MS-SomaScan translation.

For cohort integration analysis for ALS, we used the GNPC SomaScan ALS cohort containing 355 samples, including 245 ALS cases and 110 controls. These SomaScan profiles were translated into the Olink space using CKB-trained AART model and were then used as source-domain data for integration with UK Biobank Olink profiles.

### UK Biobank

UK Biobank is a population-scale prospective cohort with extensive baseline phenotyping, biological sample collection, and longitudinal linkage to health outcomes. We used the UK Biobank Pharma Proteomics Project^1^ at baseline time as the target-domain Olink dataset for ALS transfer learning. Disease diagnosis dates were collated from UKB ‘First Occurrences’ (data category 1712). ALS cases were defined using ICD-10 code (G12.2) before the baseline time. Participants whose first ALS diagnosis occurred more than 10 years before proteomic sampling were excluded, and post-proteomics ALS diagnoses were not treated as cases (*n* = 19). Controls were defined as participants with no ALS diagnosis code and were sampled at a 1:5 case-control ratio with matched age and sex. The resulting UK Biobank Olink dataset served as the target-domain cohort for ALS classification.

### Platform-specific quality controls and preprocessing

#### CKB preprocessing

For CKB, Olink NPX values and SomaScan RFU values were obtained after source study quality controls and normalization^20^. We used the non-ANML SomaScan data for the main Olink-SomaScan translation analyses, consistent with the source study comparison. SomaScan values were log-transformed before modeling. Olink and SomaScan matrices were aligned by participant identifier and UniProt-based target mapping.

#### PEX-LC preprocessing

For PEX-LC, preprocessing was performed on the 88 samples measured by both Olink and MS. The Olink matrix started from 2,923 unique Olink proteins. Individual NPX values below the assay-specific limit of detection were set to missing, and values flagged with sample-level or assay-level quality-control warnings were also set to missing. Proteins with all 88 values below the limit of detection were removed. Duplicated Olink assays mapping to the same UniProt identifier were resolved before modeling. The final Olink matrix contained 2,870 proteins.

The MS matrix started from 2,578 protein groups after removal of all-missing protein groups. Proteins with more than 50% missing values across the 88 shared samples were removed. Per-sample median centering was applied to reduce systematic TMT batch effects. The final MS matrix contained 1,747 proteins.

Remaining platform-specific missing values were imputed before model fitting. For Olink, missing values were imputed as the assay-specific limit of detection divided by ^31^. For MS, missing values were imputed using a MinProb strategy, in which low-intensity values are sampled from the lower tail of the observed protein-specific distribution. The final PEX-LC matrices contained 2,870 Olink proteins and 1,747 MS proteins. In the MS->Olink translation, 824 Olink targets had matched MS source features and 2,046 Olink targets were non-overlapping. In the Olink->MS translation, 873 MS targets had matched Olink source features and 874 MS targets were non-overlapping.

#### GNPC preprocessing

For GNPC, 983 plasma samples had matched SomaScan and MS measurements before quality controls. For SomaScan, 149 all-NaN aptamers were removed from the initial 7,745 aptamers. Local MS quality controls used protein- and sample-level call-rate filters, missingness-structure diagnostics, and outlier masking. The selected MS QC rules used a protein call-rate threshold of 0.67 and a sample call-rate threshold of 0.96. The final MS matrix contained 366 samples and 494 proteins, with 54 outlier values masked. The mean sample missingness after QCs was 1.6%.

SomaScan values were log-transformed upstream in the GNPC processing pipeline. MS values were already on a log scale after source-study quality control. Missing values and normalization for model benchmarking were handled within each cross-validation split as described below.

Target proteins with fewer than 20 finite paired observations or zero variance on either platform were excluded from per-protein evaluation. The final benchmark included 7,596 SomaScan aptamers (mapping to 6,397 unique proteins) and 494 MS proteins, with 526 overlapping and 6,066 non-overlapping proteins for the MS-to-SomaScan direction and 383 overlapping and 111 non-overlapping proteins for the SomaScan-to-MS direction.

#### Benchmark preprocessing

For benchmarking, all preprocessing steps that required estimating statistics from the data were performed within each training split and then applied to validation and held-out samples. Missing values were imputed using feature-wise medians estimated from the training split unless a platform-specific imputation strategy was defined. Features were *z*-score normalized using training-split means and standard deviations, and the same parameters were applied to validation and held-out samples. Identical preprocessing, train-validation-test splits and held-out folds were used for AART and all benchmark methods within each cohort and translation task.

### Proteomic association analysis

We performed proteomic association analyses based on 366 GNPC individuals with paired SomaScan and MS measurements for both 383 shared UniProt targets and remaining non-overlapping proteins (6,066 Somascan targets, and 111 MS targets). AART was applied in both directions, translating MS from SomaScan and SomaScan from MS. Four feature matrices were analyzed for each disease: observed SomaScan, observed MS, AART-translated SomaScan and AART-translated MS. When multiple SomaScan aptamers mapped to the same UniProt identifier, aptamer measurements were first normalized and standardized separately and then averaged to obtain one protein-level abundance value.

For each disease, we fitted a linear regression model, 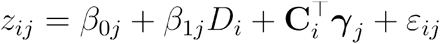, where *z*_*ij*_ is standardized protein abundance, *D*_*i*_ is the disease indicator, and *C*_*i*_ contains covariates including age and sex. The protein-specific disease effect was given by *β*_1*j*_. We compared observed-protein and translated-protein association analyses using Pearson correlation of effect sizes, agreement of effect directions, and overlap of nominally significant disease markers defined by *P* < 0.05.

### ALS classification analysis

To integrate GNPC with UK Biobank proteomic data, we first trained a SomaScan->Olink AART model using the paired CKB Olink-SomaScan cohort. The trained AART model was applied to the GNPC ALS SomaScan cohort, yielding 355 AART-translated Olink profiles (245 cases) with 2,168 proteins. UK Biobank ALS cases were defined as described above. For each of five random seeds, we used all UK Biobank ALS cases (*n* = 19) and 95 randomly-sampled matched controls, split them by 80/20 into UKB-train (80%) and test sets (20%), and progressively added a fraction *f* ∈{0%, 10%, 25%, 50%, 75%,100%} of GNPC-translated samples. Within-cohort feature standardization was performed before integrating UKB and GNPC-translated profiles. For each random seed and augmentation fraction, proteins were first selected in the combined training set using covariate-adjusted abundance regression analysis with Benjamini–Hochberg FDR < 0.05, and then a class-weight-balanced -regularized logistic regression classifier was fitted on the pre-selected proteins and evaluated by AUROC on the held-out UKB test set.

To benchmark against existing ALS biomarkers, we used an established 17-protein reference panel^8^. After intersecting this panel with the SomaScan–Olink overlapping protein set, 12 proteins remained (NEFL, ART3, MEGF10, CSRP3, CA3, EDA2R, MYOM3, ACTN2, HS6ST2, KLK4, HSPB6, and GZMH). We recorded the selection frequency of each protein across random seeds and settings to assess reproducibility.

### Gene function analysis

To characterize the biological properties of targets accurately translated by AART, we performed functional enrichment analysis using g:Profiler^49^. Enrichment was tested against Gene Ontology Biological Process, Molecular Function and Cellular Component terms, Human Protein Atlas tissue annotations, and Reactome pathways. Analyses were performed separately for each translation direction using a direction-specific custom background consisting of all unique genes corresponding to target features evaluated in that task. This background was used to reduce enrichment artifacts caused by platform-specific assay coverage. Statistical significance was assessed using the g:SCS multiple-testing correction, with FDR < 0.05 considered significant. In enrichment testing, assay features were first mapped to gene identifiers. If multiple features mapped to the same gene, that gene was counted only once in both the query set and the background.

## Extended Data Figures

**Extended Data Fig. 1.**
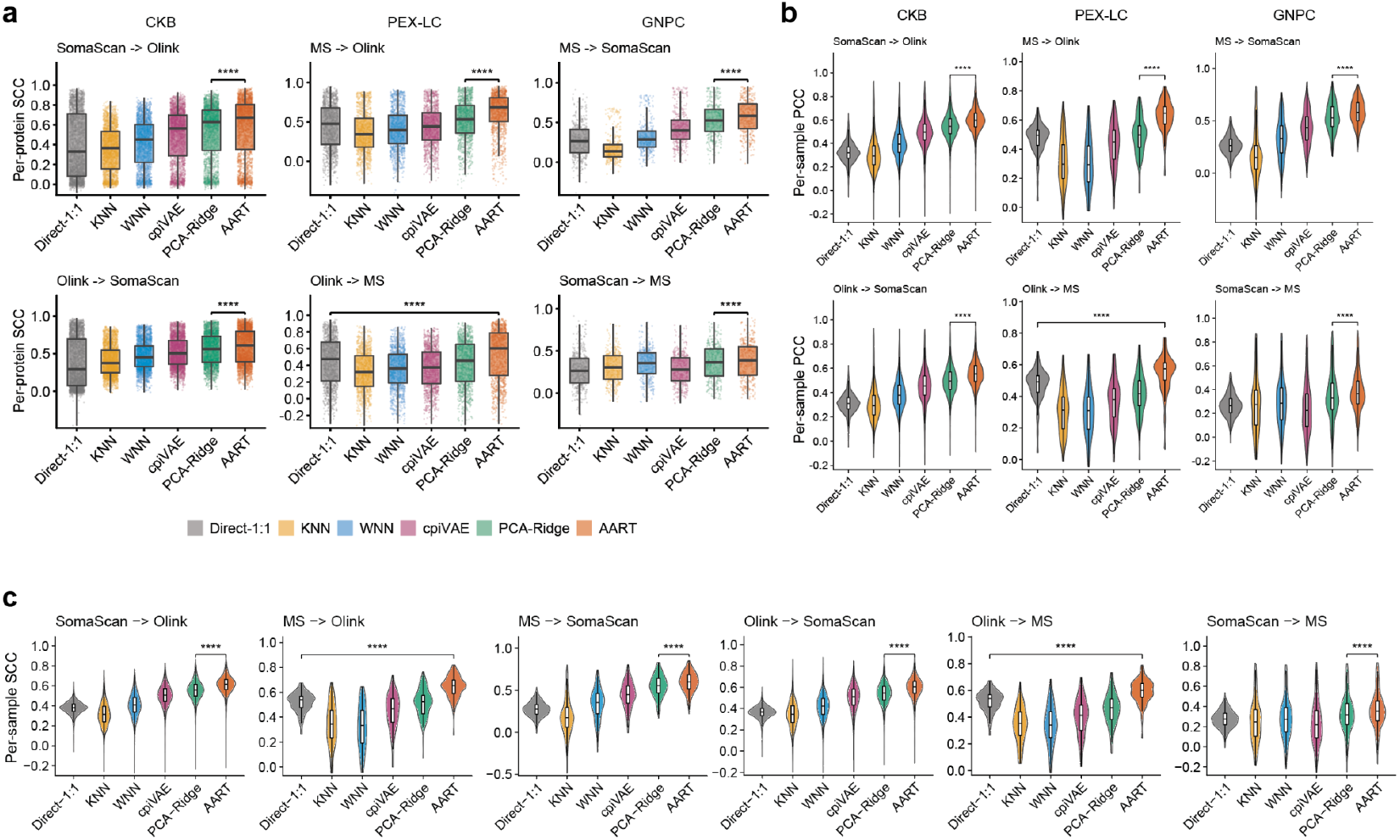
Translation performance comparison using alternative metrics. **a**, Box plots show five-fold cross-validated, per-protein Spearman correlations between translated and observed values for overlapping target proteins across six translation tasks: SomaScan->Olink and Olink->SomaScan in CKB (*n* = 3,976), MS->Olink and Olink->MS in PEX-LC (*n* = 88), and MS->SomaScan and SomaScan->MS in GNPC (*n* = 366). The box plot center line, limits, and whiskers represent the median, quartiles, and 1.5x interquartile range (IQR), respectively. ****, *P* < 1e-4 by one-sided paired Wilcoxon signed-rank test. SCC, Spearman correlation coefficient. **b**, Violin plots show per-sample Pearson correlations across target proteins. ****, *P* < 1e-4 by one-sided paired Wilcoxon signed-rank test. PCC, Pearson correlation coefficient. **c**, Violin plots show per-sample Spearman correlations across target proteins. ****, *P* < 1e-4 by one-sided paired Wilcoxon signed-rank test. All comparisons were performed between AART and the best baseline method. Identical five-fold training, validation, and test splits were used across methods within each task. Per-protein correlations were calculated across test samples; per-sample correlations were calculated across target proteins.

**Extended Data Fig. 2.**
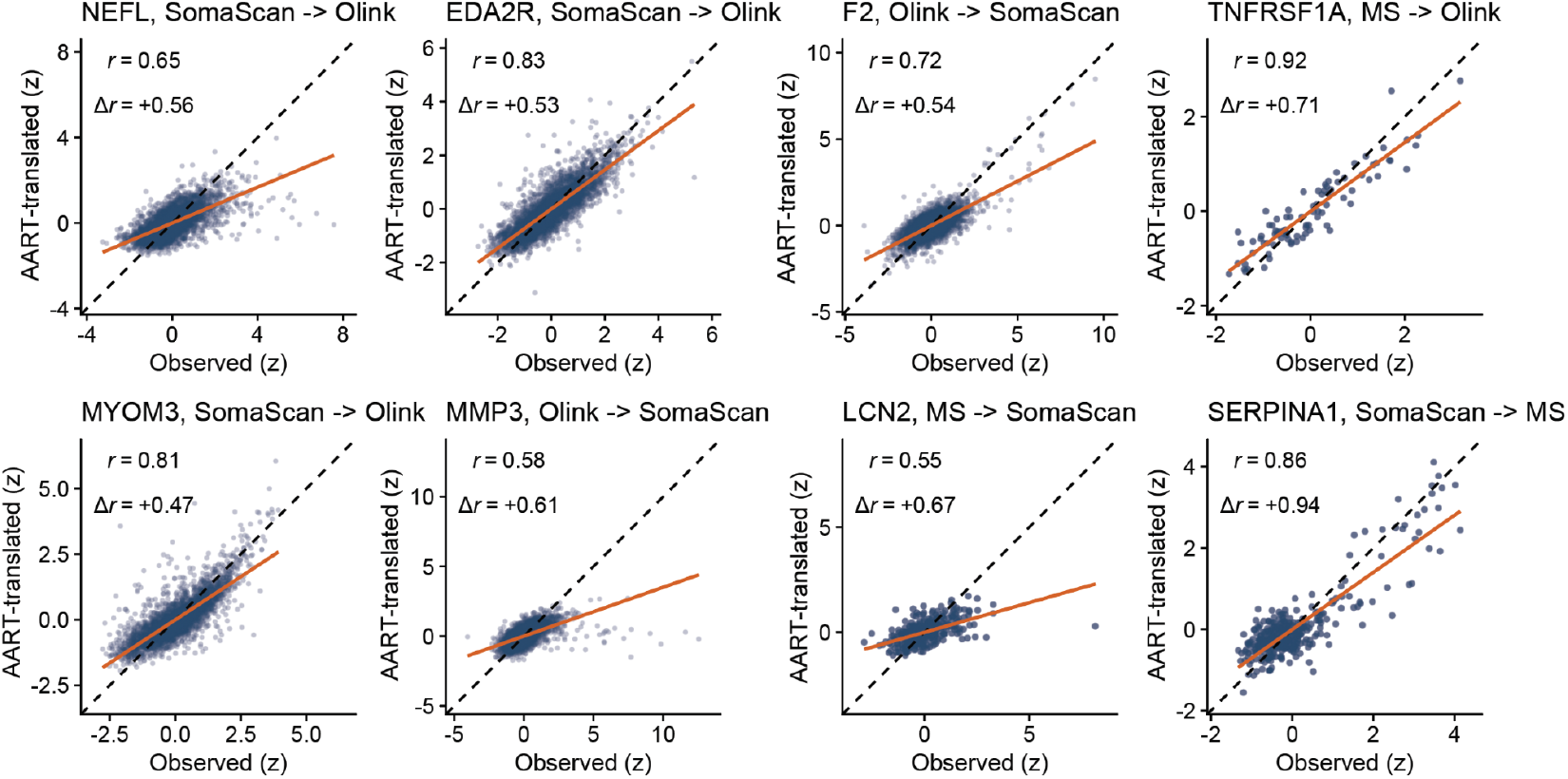
Representative protein-level translation results. Scatter plots show AART-translated versus observed *z*-score normalized protein abundance in held-out samples for eight overlapping target proteins: NEFL, EDA2R, and MYOM3 for CKB SomaScan->Olink; F2 and MMP3 for CKB Olink->SomaScan; TNFRSF1A for PEX-LC MS->Olink; LCN2 for GNPC MS->SomaScan; SERPINA1 for GNPC SomaScan->MS. Each dot represents one held-out test sample. Orange lines indicate the ordinary least-squares fits. *r* denotes Pearson correlation coefficient calculated between observed and AART-translated abundances, and Δ*r* denotes the difference between AART- and Direct-1:1-derived correlations.

**Extended Data Fig. 3.**
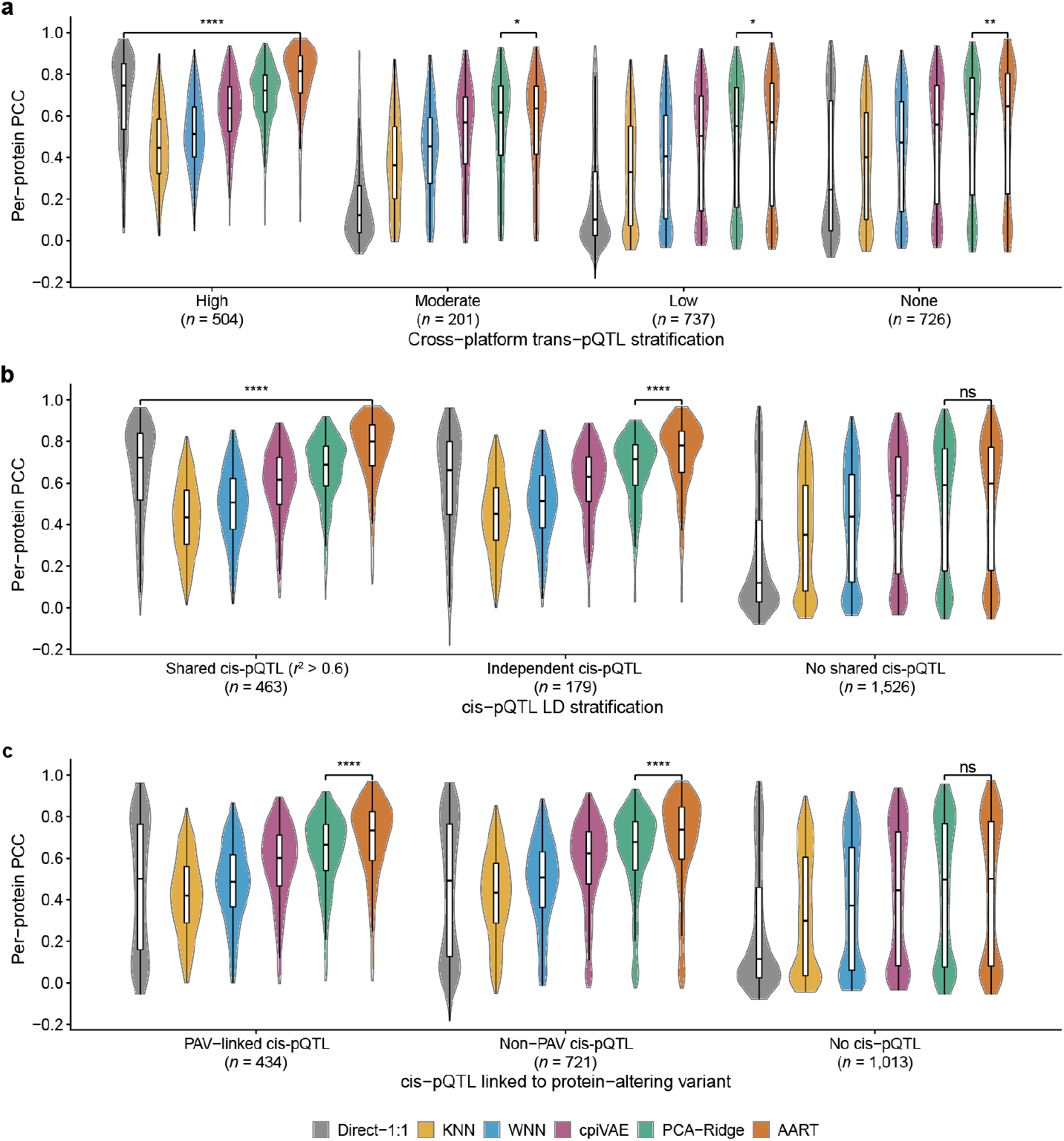
Genetic stratification of CKB SomaScan->Olink translation accuracy. **a-c**, Violin plots show five-fold cross-validated per-protein Pearson correlations for CKB SomaScan->Olink translation, stratified by genetic concordance annotations including trans-eQTL (**a**), cis-eQTL LD (**b**), and cis-eQTL linked to protein-altering variants (**c**). In **a**, rows stratify target proteins by cross-platform trans-pQTL concordance (High, proteins with colocalized trans-pQTLs across platforms; Moderate, proteins with trans-pQTLs detected on both platforms without colocalization; Low, proteins with trans-pQTL detected on one platform only; None, proteins with no trans-pQTL detected on either platform). In **b**, rows stratify proteins by linkage disequilibrium (LD) between cross-platform cis-pQTL signals, in which “shared cis-eQTL” is defined by at least one cis-pQTL pair with LD *r*^2^ > 0.6. In **c**, rows stratify proteins by cis-pQTLs linked to protein-altering variants (PAVs). The box plot center line, limits, and whiskers represent the median, quartiles, and 1.5x interquartile range (IQR), respectively. *P*-values by one-sided paired Wilcoxon signed-rank tests. Comparisons were performed between AART and the best baseline method. *, *P* < 0.05; **, *P* < 0.01; ****, *P* < 0.0001; ns, not significant; PCC, Pearson correlation coefficient.

**Extended Data Fig. 4.**
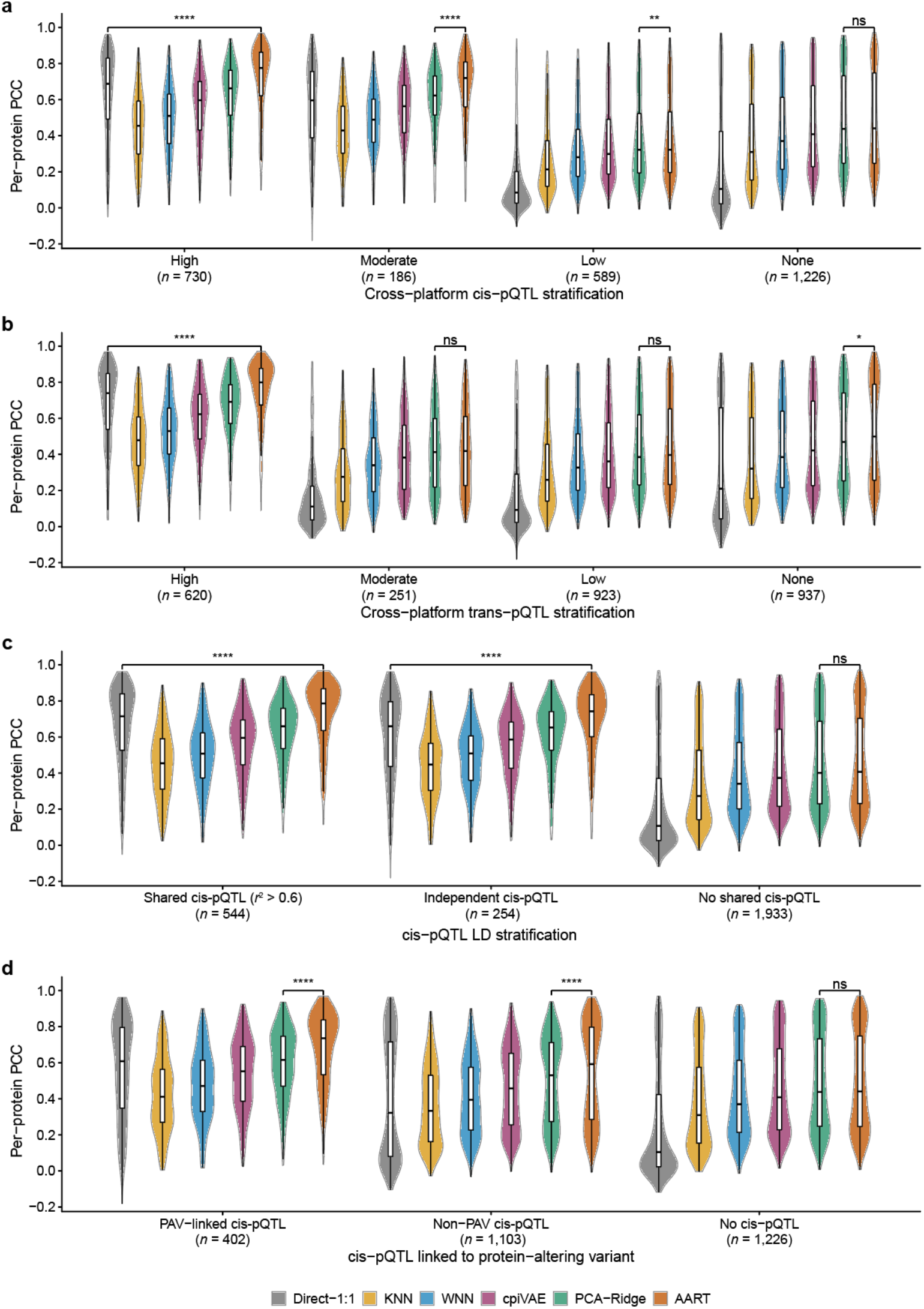
Genetic stratification of CKB Olink->SomaScan translation accuracy. **a-d**, Violin plots show cross-validated per-protein Pearson correlations for CKB SomaScan->Olink translation, stratified by genetic concordance annotations including cis-eQTL (**a**), trans-eQTL (**b**), cis-eQTL LD (**c**), and cis-eQTL linked to protein-altering variants (**d**). In **a**, rows stratify target proteins by cross-platform cis-pQTL concordance (High, proteins with colocalized cis-pQTLs across platforms; Moderate, proteins with cis-pQTLs detected on both platforms without colocalization; Low, proteins with cis-pQTL detected on one platform only; None, proteins with no cis-pQTL detected on either platform. In **b**, rows stratify target proteins by cross-platform trans-pQTL concordance (High, proteins with colocalized trans-pQTLs across platforms; Moderate, proteins with trans-pQTLs detected on both platforms without colocalization; Low, proteins with trans-pQTL detected on one platform only; None, proteins with no trans-pQTL detected on either platform). In **c**, rows stratify proteins by linkage disequilibrium (LD) between cross-platform cis-pQTL signals, in which “shared cis-eQTL” is defined by at least one cis-pQTL pair with LD *r*^2^ > 0.6. In **d**, rows stratify proteins by cis-pQTLs linked to protein-altering variants (PAVs). The box plot center line, limits, and whiskers represent the median, quartiles, and 1.5x interquartile range (IQR), respectively. *P*-values by one-sided paired Wilcoxon signed-rank tests. Comparisons were performed between AART and the best baseline method. *, *P* < 0.05; **, *P* < 0.01; ****, *P* < 0.0001; ns, not significant; PCC, Pearson correlation coefficient.

**Extended Data Fig. 5.**
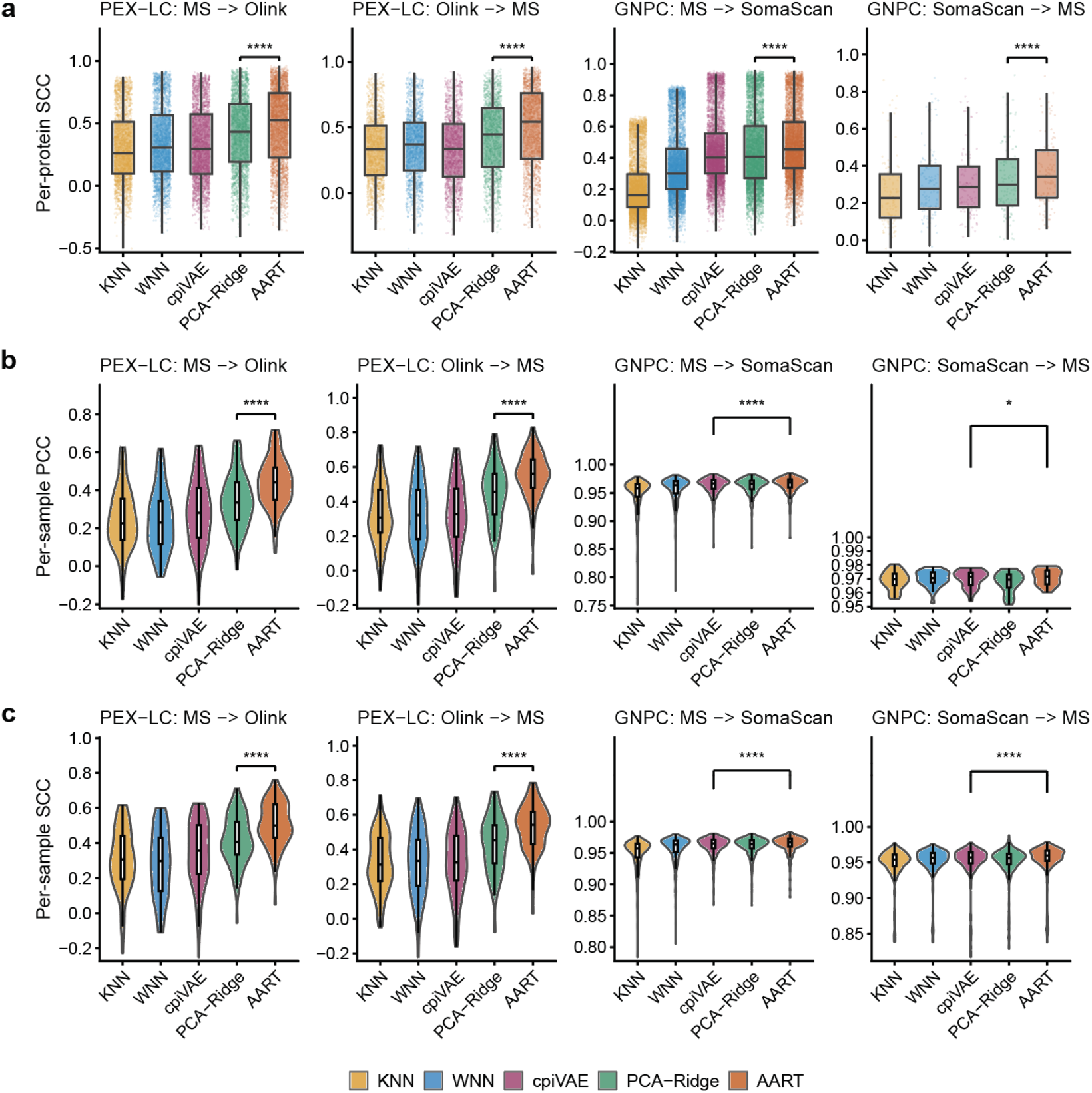
Translation performance for non-overlapping targets using alternative metrics. **a**, Box plots show five-fold cross-validated, per-protein Spearman correlations between translated and observed values for non-overlapping target proteins across four translation tasks: MS->Olink and Olink->MS in PEX-LC (*n* = 88), and MS->SomaScan and SomaScan->MS in GNPC (*n* = 366). SCC, Spearman correlation coefficient. **b**, Violin plots show per-sample Pearson correlations across target proteins. PCC, Pearson correlation coefficient. **c**, Violin plots show per-sample Spearman correlations across target proteins. The box plot center line, limits, and whiskers represent the median, quartiles, and 1.5x interquartile range (IQR), respectively. Comparisons were performed between AART and the best baseline method. *P*-values by one-sided paired Wilcoxon signed-rank tests. *, *P* < 0.0s5; ****, *P* < 0.0001. Identical five-fold training, validation, and test splits were used across methods within each task. Per-protein correlations were calculated across test samples; per-sample correlations were calculated across target proteins.

**Extended Data Fig. 6.**
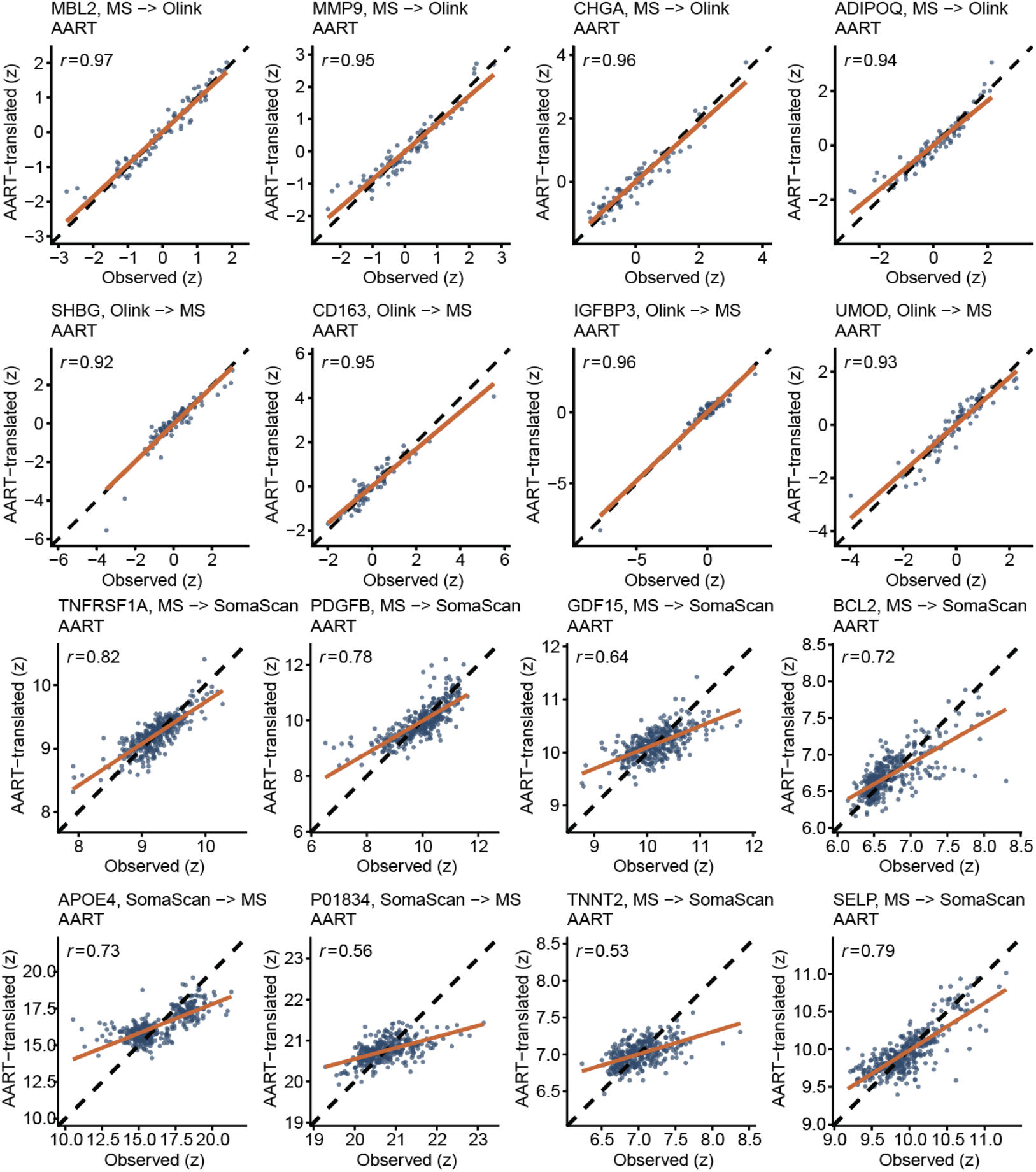
AART translation results for representative non-overlapping proteins. Scatter plots show AART-translated versus observed *z*-score normalized protein abundances across held-out samples for 16 non-overlapping proteins: MBL2, MMP9, CHGA, and ADIPOQ for PEX-LC MS->Olink; SHBHG, CD163, IGFBP3, and UMOD for PEX-LC Olink->MS; TNFRSF1A, PDGFB, GDF15, and BCL2 for GNPC MS->SomaScan; APOE4, P01834, TNNT2, and SELP for GNPC SomaScan->MS. Each dot represents one held-out test sample. Orange lines indicate the ordinary least-squares fits. *r* denotes Pearson correlation coefficient calculated between observed and AART-translated abundances.

**Extended Data Fig. 7.**
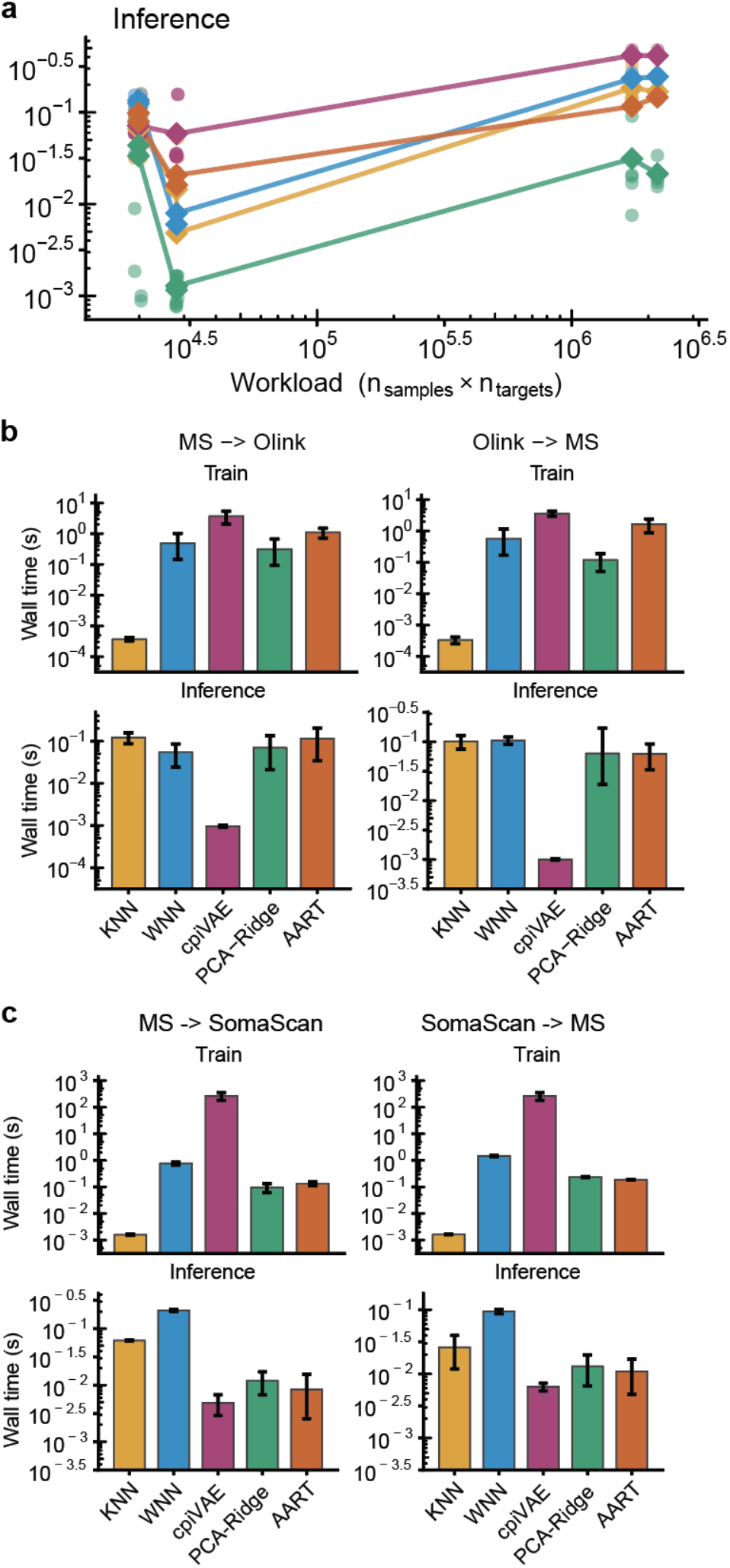
Runtimes for AART and baseline methods. **a**, Inference wall time plotted against workloads, defined as the product of sample size and number of target features. Round dots denote individual runs and square dots denote method-level means. **b-c**, Training and inference wall time for PEX-LC MS-Olink (**b**) and GNPC MS-SomaScan non-overlapping protein translation (**c**). Wall time is shown on a log scale. Bars represent mean wall time across folds and repeated runs, and error bars denote standard errors.

**Extended Data Fig. 8.**
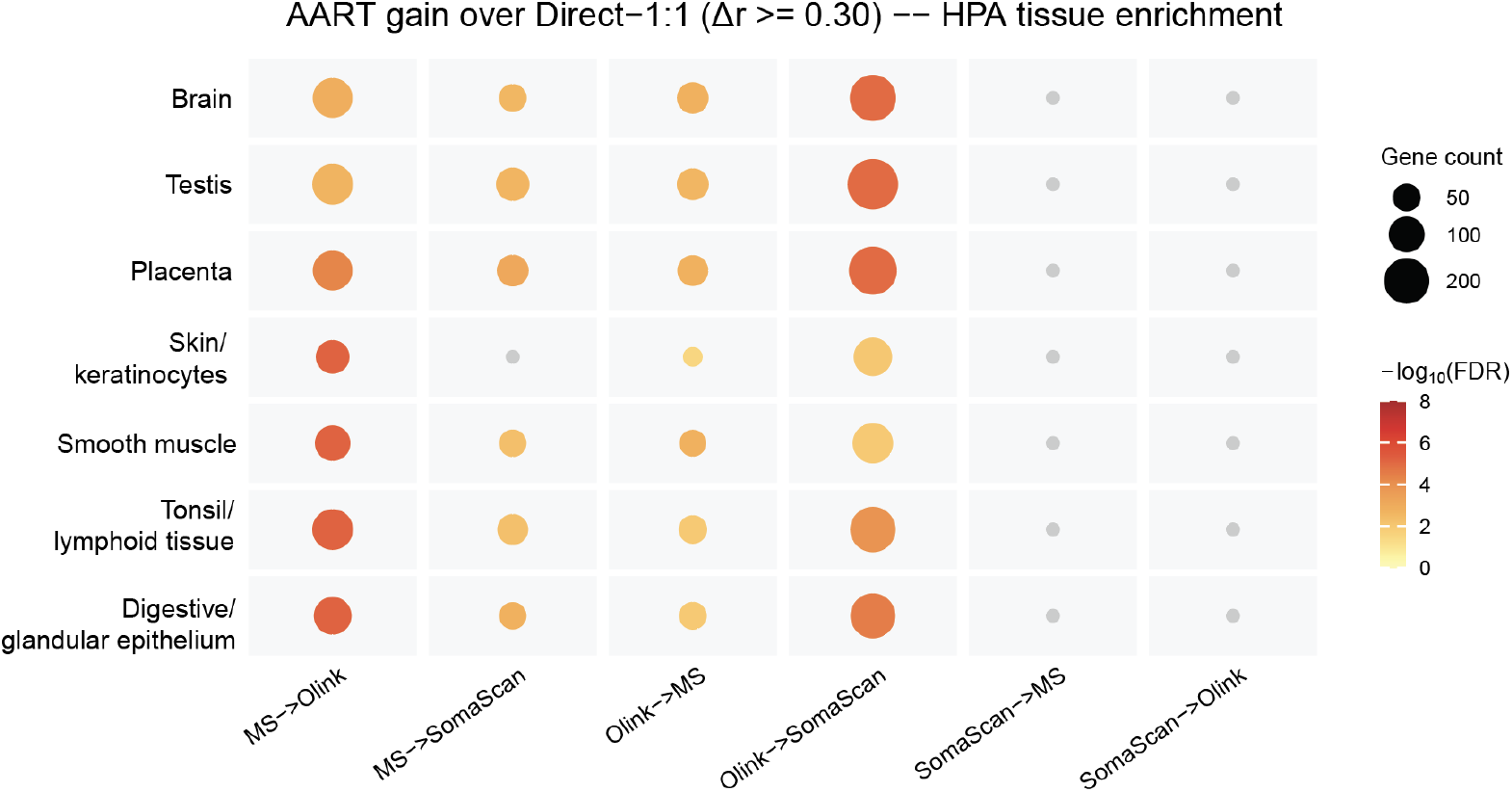
Gene function enrichment of AART-better-translated overlapping proteins. Human Protein Atlas (HPA) tissue enrichment of overlapping proteins whose translation was substantially improved by AART compared with Direct-1:1 (Δ*r* >= 0.30). Bubble color denotes −log_10_(FDR), grey bubble represents non-significance, and bubble size indicates number of genes. MS, mass spectrometry.

**Extended Data Fig. 9.**
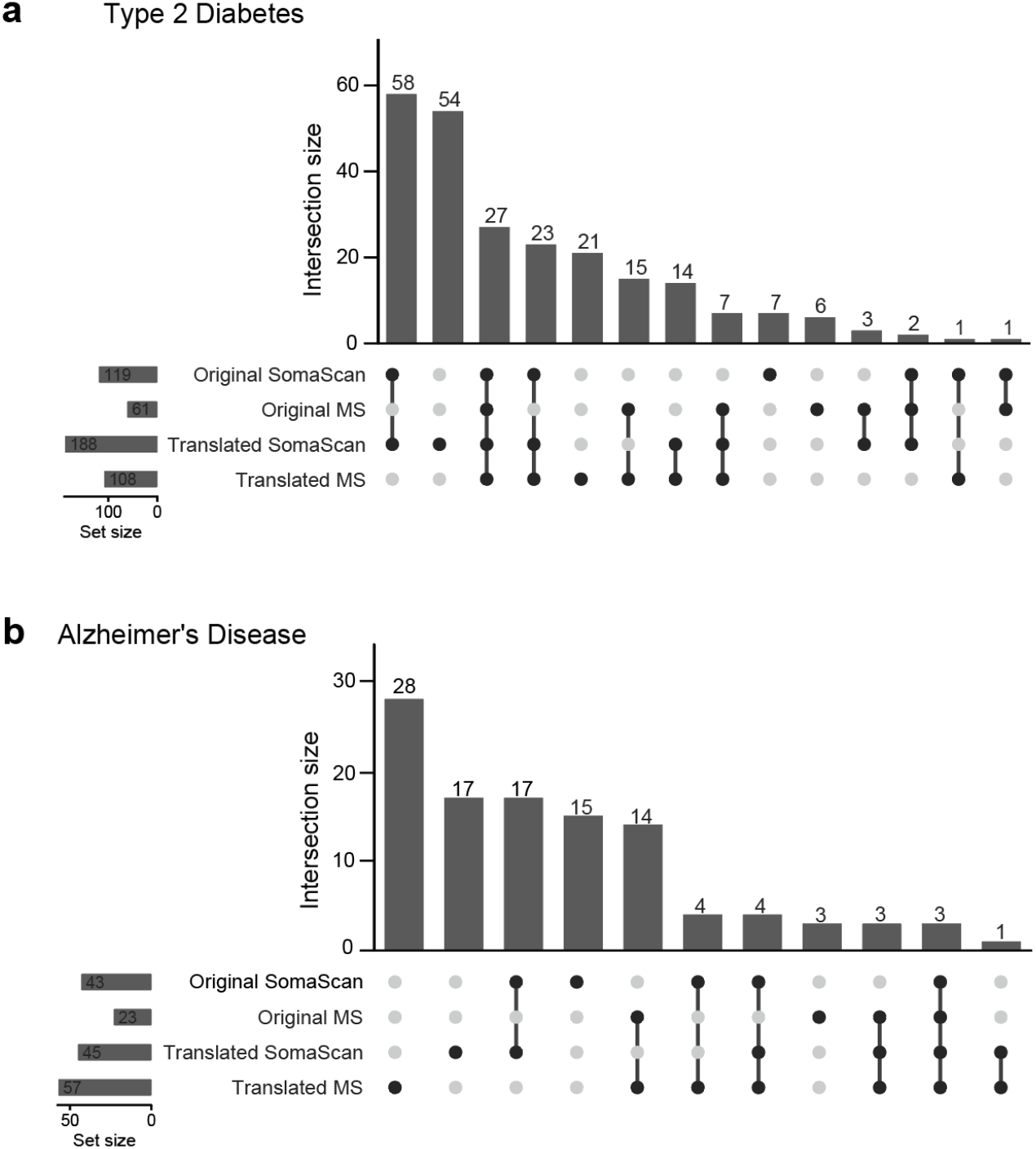
Overlaps of disease-associated proteins identified based on observed and AART-translated overlapping proteins. **a-b**, UpSet plots show sets of nominally significant disease-associated proteins among the 383 overlapping proteins in the paired GNPC SomaScan-MS subset (*n* = 366 samples) for type 2 diabetes (**a**) and Alzheimer’s disease (**b**), respectively. Proteomic association analyses were performed separately based on observed SomaScan, observed MS, AART-translated SomaScan, and AART-translated MS measurements. Nominal significance was defined as *P* < 0.05. Horizontal bars denote the number of significant proteins in each analysis, and vertical bars show intersection sizes.

**Extended Data Fig. 10.**
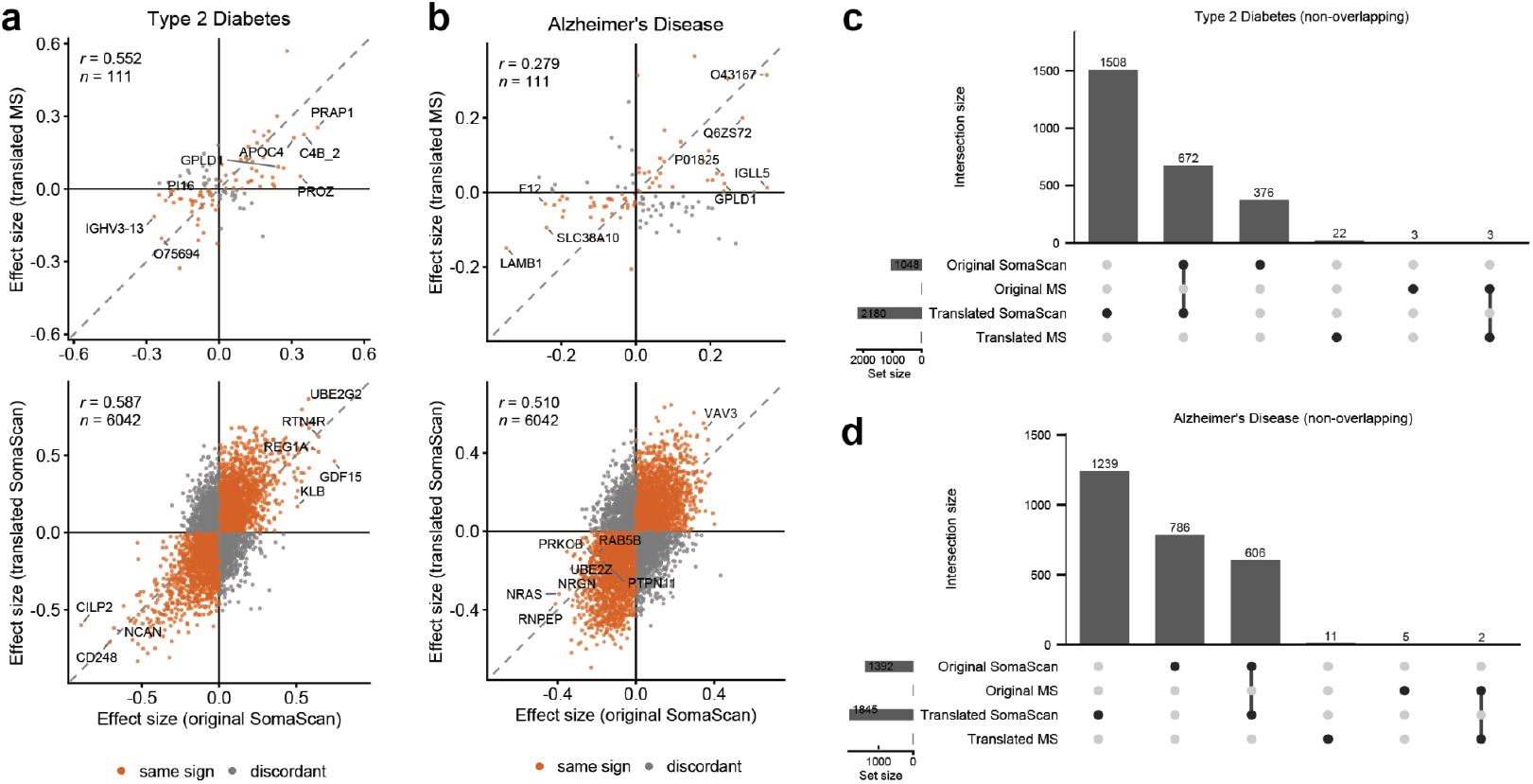
Proteomic association analyses for non-overlapping proteins. **a-b**, Protein associated effects estimated based on observed (x-axis) and AART-translated non-overlapping proteins (y-axis) for type 2 diabetes (**a**) and Alzheimer’s disease (**b**), respectively. Each dot represents one protein; orange dots denote concordant effect directions and grey dots denote discordant directions. Representative proteins with large absolute effects are highlighted. **c-d**, UpSet plots show sets of nominally significant non-overlapping target features for type 2 diabetes (**c**) and Alzheimer’s disease (**d**) in observed and AART-translated SomaScan and MS analyses. Nominal significance was defined as *P* < 0.05. Horizontal bars denote set sizes and vertical bars show intersection sizes. All analyses were performed using the SomaScan-MS paired GNPC subset (*n* = 366) with 6,066 non-overlapping somascan proteins and 111 non-overlapping MS proteins.

